# Kinetic windows for abiogenesis

**DOI:** 10.64898/2026.11.20.768022

**Authors:** Stephen J. Mojzsis, Anna Medvegy, Barbara Kremer

## Abstract

The origin of life on Earth occurred within a narrow window delimited by cessation of intense late accretion bombardment at ca. 4.4 Ga on the one hand, and younger (<4.2 Ga) evidence gleaned via molecular phylogenetic studies, isotopic biosignatures and morphological traces on the other. Molecular clock analyses tentatively place the median age for the Last Universal Common Ancestor (LUCA) population of all contemporary DNA life at ca. 4.2 Ga. Cryptic isotopic biosignatures go back as far as 4.1± 0.01 Ga in Hadean zircons (δ¹³C = -24 ± 5‰) and more robustly in Eoarchean rocks at ca. 3.83 Ga (δ¹³C ≈ -35‰). Morphological remains including microfossils and microbially mediated sedimentary structures are almost as ancient (ca. 3.5 Ga). To make better sense of this early and rapid timeline and its implications for a biosphere on Hadean Earth, we parameterize the *kinetic windows* for that transition from non-living to living chemistry (i.e. abiogenesis). Here, we reinforce the finding that before the LUCA, a RiboNucleic Acid (RNA) and/or RiboNucleoProtein (RNP) World could have persisted for <130 Myr. Furthermore, we show that conditions which define early Earth’s kinetic windows stand out compared to other worlds in our Solar System considered as candidates for abiogenesis. Specifically, these are sustained stable liquid water from ca. 4.4 Ga, wet-dry cycling on dry land under an atmosphere, effective crustal recycling to rejuvenate disequilibrium chemistry, available key elements for synthesis and growth, mechanisms to concentrate prebiotic reaction products without also destroying them, and sufficient spatial scales for subsequent evolution and diversification. We argue that the steps from abiogenesis to DNA required chemistry coordinated with environmental conditions overlapping in time with each transition – from prebiotic synthesis to cellular life – governed by distinct rate-limiting processes operating over geologically short (10^0^–10⁸ y) timescales. Our analysis suggests that if DNA life had not emerged, an informationally deficient and thermally unstable RNA biosphere would eventually have gone extinct.

## 1. Introduction

Nearly every aspect of the origin of life is debated. This situation arises because the physical, chemical and temporal bounds for the transition from non-living matter to life (e.g. **Dana, 1866**), conventionally termed abiogenesis, are obscured or erased by all subsequent geological and biological events that have taken place on our planet. Earth effectively destroys its own vestiges of a beginning (**Hutton, 1788**). As for what is confidently preserved in terms of the early record of life, a relatively sophisticated microbial biosphere at the start of the Paleoarchean era ca. 3.5 Ga is well attested thanks to morphological evidence from interpretable microfossils and biologically mediated sedimentary structures such as stromatolites (e.g. **Walter et al., 1980; Schopf, 1993; Allwood et al., 2009; Kremer and Kaźmierczak, 2017**). Our focus here, however, is on times earlier than this, within the roughly billion-year interval between planet formation and the oldest (albeit contested) biosignatures. Our greatest challenge to understanding this formative time is that while it witnessed a complex sequence of prebiotic chemistry, molecular self-organization, and evolutionary innovation, life’s emergence occurred in environments for which no direct biological records are preserved and scant geochemical records survive. How then can we parameterize the processes responsible for abiogenesis? We argue that effective parameterization requires going beyond cataloguing the various chemical ingredients of life and documenting the reaction products in various prebiotically plausible settings. Instead, it makes sense to quantify the *kinetic windows* – defined here as the temporal intervals during which specific conditions permitted particular stages of abiogenesis to proceed and persist – to understand why life emerged on Earth very early and apparently not elsewhere in the Solar System. The outcome of our analysis consistently yields quantitatively short timelines for abiogenesis and biocompatibility. Biocompatibility is defined here as those necessary physical and chemical conditions which must be in place to allow life to exist on the surface of a planet, regardless of whether or not life is present (e.g. **Fogg, 1992**).

Our motivation to define kinetic windows stems from the many advances in biochemistry, geochemistry and cosmochemistry which have fundamentally transformed our understanding of the nature of abiogenesis. For one, progress in RNA chemistry and prebiotic syntheses have quantified the environmental sensitivities and rate-limiting steps important for the early biochemical pathways toward an RNA World (e.g. **Powner et al., 2009; Sutherland et al., 2016; Krishnamurthy et al., 2017; Benner et al., 2020**). Advances in bioinformatics have also revolutionized our understanding. Molecular clock analyses calibrated with pre-LUCA gene duplications place the Last Universal Common Ancestor (LUCA) population of all contemporary DNA life at around 4.2 Ga, albeit with large uncertainties (**Feng et al., 1997; Battistuzzi et al., 2004; Moody et al., 2024**). Intrinsic Hadean (pre-4 Ga) records of crustal processes, interactions with liquid water, redox conditions for melting of Earth’s mantle and plate-boundary scenarios have been revealed by detailed geochemical analysis of the oldest terrestrial zircons (e.g. **Mojzsis et al., 2001; Trail et al., 2011; Harrison, 2009**). For instance, high-precision geochronology combined with stable isotopic analysis of Hadean zircon inclusions have pushed back tantalizing hints of a potential biosignature back to 4.1 Ga (**Bell et al 2015**). The oldest actual rocks of sedimentary origin are nearly as old as this, at ca. 3.83 Ga, and host isotopic signatures interpreted to record biological activity in a marine setting (**Mojzsis et al., 1996; Long et al., 2025**). Important extrinsic effects to the Hadean environment have also come into focus. Refined models for the nature and timing of late accretion bombardments are now used to constrain the onset of continuous biocompatible conditions on Hadean Earth (**Abramov and Mojzsis, 2009; Mojzsis et al., 2019; Abramov et al. 2026**). As transformative as these discoveries are in supporting the great antiquity of the biosphere, they remain at odds with the prevailing view that the environment of Hadean Earth could not support the origin of life. For instance, it has long been argued that if a precursor RNA World stage actually existed (**Rich, 1962; Crick, 1968**) it did so as part of the process of the origin of life and as a *lengthy* intermediate stage on the path DNA, perhaps spanning half a billion years or more (**Joyce, 2002**).

Collectively, the findings cited above lead us to converge on a striking conclusion: The path from abiogenesis to the RNA- and/or RNP World (e.g. **Gilbert, 1986; Müller et al., 2022**) occurred over a relatively short time in the middle part of the Hadean eon (**Abramov et al., 2026**). We interpret this compressed timeline as a reflection of the kinetic constraints fundamentally imposed by RNA’s intrinsic biochemical instability in the geochemical Hadean environment, such as its susceptibility to alkaline hydrolysis and inability to form stable hydrophobic cores (**Keller et al., 2017; Forster and Walde, 2022**). This realization in turn leads us to consider that the transition to DNA-based genomes was not merely an evolutionary refinement but a kinetic necessity (e.g. **Walton et al., 2026**). Our conclusion is that the transition to DNA was driven by a combination of pressures which challenged the severely limited achievable inherent complexity of a postuled RNA/RNP organism.

The kinetic windows concept frames the key constraints to understanding what led to the inevitable decline of the RNA/RNP World while paving the way for the DNA World and the LUCA. Unlike thermodynamic favorability, which determines whether a reaction can occur, kinetic accessibility determines whether it will occur on geologically relevant timescales and within natural constraints. Multiple kinetic windows have to align for abiogenesis in a biocompatible environment: Synthesis of nucleotide precursors and their effective polymerization into functional oligomers in the emergence of self-replicating systems, the transition to cellular organization and the establishment of metabolism all leading to a living entity. The conceptual framework of Gánti (**1997, 2003**) is useful here because it effectively defines the minimal living entity (chemoton) as a chemically coupled triad consisting of an autocatalytic metabolic network, a template replicating genetic polymer, and an encapsulating membrane in close alignment with modern working definitions of a biological system (**Joyce, 1994**).

Each kinetic window is defined by specific physical-chemical parameter – temperature, pH, water activity, UV flux, and elemental concentrations – that must fall within narrow ranges for extended periods (10^0^-10^8^ years) to allow the first life to emerge and persist long enough to evolve to DNA. First, we establish the geological framework that constrains when abiogenesis could have occurred, then develop a formal definition of kinetic windows applicable to abiogenesis scenarios. Next, we examine specific constraints: the physical-chemical parameter space (temperature, pH, radiation, water activity), the duration and limitations of the RNA World, the possible biogeodynamic triggers for its extinction, the nature of LUCA and the DNA transition, and touch on the critical role of nutrient availability. Then, we compare Earth’s kinetic windows with those of other Solar System bodies for biocompatibility, including the early warm interiors of carbonaceous asteroids and the cloaked oceans of icy moons of the gas giants and the distant Trans-Neptunian Objects of the outer Solar System. We conclude by exploring implications of the approach for the probability and timescale of abiogenesis and identifying key uncertainties requiring future investigation.

## 2. The Lower Kinetic Bookend – Planet Formation and Early Bombardment

Earth’s formation was nearly complete by 4.54 Ga through accretion of building blocks mostly from its local feeding zone (e.g. **Brasser et al. 2018**), perhaps in a disk sub-structure such as a ring or pressure bump (**Brasser and Mojzsis, 2020**). This primary episode of formation was followed soon afterwards by the Moon-forming impact of a roughly Mars-sized planetary embryo named *Theia* (which was at least 10% of Earth’s mass) at 4.51 ± 0.01 Ga (**Barboni et al., 2017; Canup, 2004; Halliday, 2000**). If there was any pre-existing life or proto-life on our embryonic planet at that time, the Moon-forming event effectively sterilized it. Moon formation therefore sets a hard boundary for any abiogenesis proposed timelines (**cf. Sleep et al., 2001**). In the aftermath of the Moon another colossal impact event is postulated to have occurred. Dubbed the Late Veneer, this collision may have augmented Earth’s mass by approximately 0.5-1 wt.% with an object dubbed *Moneta* (**Genda et al. 2017**). It has been argued that such a relatively late (4.48-4.45 Ga) impact of a lunar mass object (2700 km diameter; **Bottke et al. 2015; Brasser et al., 2016**) can account for the highly siderophile element abundance in chondritic relative proportions of Earth’s mantle (e.g. **Walker, 2009**). If the Late Veneer happened, and this too is debated (**cf. Wang et al., 2023**), it further delimits the kinetic window for life’s emergence as having taken place sometime after ca. 4.45 Ga. No other similarly immense impact with the capacity to sterilize the planet has affected Earth since *Moneta* time (**Abramov et al., 2026**).

Dynamical studies of planet formation coupled with the geochemistry and cosmochemistry have revised much of what we knew about the history of late accretion and thus, the thermal properties of the Hadean Earth surface. For instance, giant planet migration, previously postulated to have triggered a “late heavy bombardment” (LHB) at ∼3.9 Ga, is now understood to have commenced much earlier, at or before ∼4.45 Ga, based on the absence of wholesale crustal reset ages in highly retentive chronometers (e.g. U-Pb and Pb-Pb in zircon and apatite) from some asteroidal meteorites (e.g. **Walton et al., 2026**). This migration triggered an intense but short-lived (∼30 Myr) cometary influx beginning at 4.48 Ga and ending by 4.45 Ga, which included accretion of leftover planetesimals and asteroids (**Brasser et al., 2020, 2021**). The sequential termination of radiometric age resets in the meteorite data as a function of differential closure temperatures (**Dodson, 1973**) of mineral systems provides a quantitative record of this declining impact flux (**Mojzsis et al., 2019**). Closure temperature of a geochronological system can be defined as its temperature at the time corresponding to its apparent age and not necessarily its crystallization age. This is important, because highly retentive U-Pb and Pb-Pb geochronology (T*_c_* ∼1000°C) studies show age resets ending very early at ∼4.45 Ga, while intermediate-closure-temperature systems (whole-rock Pb-Pb and Sm-Nd, Tc ∼600°C) terminate around 4.40 Ga. This thermal timeline from the geochronology is in line with that predicted by the planetary dynamic models cited above. Less retentive ⁴⁰Ar-³⁹Ar ages pick up near 4.48 Ga and show a continuum extending from ∼4.48 to 3.0 Ga, indicating a long monotonic decline of leftover debris from accretion rather than an LHB spike. It is important to emphasize that these various models show that the population of large impactor masses needed to bring about global-scale crustal melting and sterilization (objects ≥9.31 × 10²¹ kg) effectively ran out by ∼4.45 Ga (e.g. **Abramov et al. 2026**). Three-dimensional thermal bombardment models also show that the abating impact flux after ∼4.4 Ga was insufficient to sterilize the surface zone, defined as temperatures ≥130°C for the upper 4 km of crust (**Abramov and Mojzsis, 2009**). This in turn implies that some relatively low temperature niches were established for prebiotic chemistry (**Abramov et al., 2026**) within approximately 170 Myr after the first Solar System solids formed (e.g. **Bouvier and Wadhwa, 2010**). Besides the Hadean terrestrial zircon record cited above, the survival of martian zircons at 4.428–4.470 Ga (**Humayun et al., 2013**) and lunar zircons at 4.47 Ga (**Greer et al., 2023**) provides separate proof that rocky crusts of other worlds in the inner Solar System did not experience wholesale melting after about 4.45 Ga, otherwise these zircon grains would not exist. After the epoch of colossal impacts came to an end, the inner Solar System was primed for life by the middle part of the Hadean eon.

### 2.1 Intrinsic Hadean Biocompatibility: Hydrosphere and Atmosphere

The presence of liquid water is a prerequisite for the biochemistry that we know, and multiple lines of evidence indicate that Earth possessed a stable hydrosphere by about 4.4 Ga or within 160 Myr of Solar System formation (e.g. **Miyazaki and Korenaga, 2022**). The existence of a hydrosphere necessarily implies the presence of a substantial atmosphere to provide the pressure and temperature conditions allowing for stable liquid water at the surface. It is important to note that the record of such a liquid water hydrosphere – as opposed to supercritical fluids – is implied by the stable isotope fractionation of oxygen (^18^O/^16^O) in the clay-contaminated melt source of the Hadean zircons (**Mojzsis, 2026; cf. Kasting and Ono, 2006**).

Zircon crystallization temperatures, stable isotopic values, trace element compositions and inclusion assemblages show that Hadean crust interacted with liquid water (**Harrison, 2009**), but what does this really mean for the nature of the surface and abiogenesis? Geodynamic models coupled with geochemical data show that the Hadean crust was affected by intrinsic processes including local or transient subduction which variably recycled the primordial crust leading to more evolved silica-rich melts which in turn give rise to buoyant crust (**Gerya et al., 2015; Caro et al., 2025**). Other evidence for non-marine surface environments has come to light form the documentation of exceptionally low ¹⁸O/¹⁶O values recorded in some Hadean zircons pointing to meteoric (rain) water-crust interactions (**Gameleldian et al., 2024**). Most commonly, these signatures can be interpreted to mean that at least groundwater – and very likely lakes and even seas – existed before 4 Gyr ago on exposed landmasses collecting rainfall into catchment basins. These global conditions eminently allow for wet-dry cycling in shallow ponds or hydrothermal systems, necessary prerequisite to many prebiotic chemistry scenarios (**e.g. Benner et al., 2010; Damer and Deamer, 2015; Sutherland, 2016**). This is important because remelting of crust provides disequilibrium chemistry, wet-dry cycles concentrate reactants and promote polymerization reactions, while provide mineral surfaces for catalysis (**Kelley et al., 2001; Deamer and Dworkin, 2005**).

Precise details about atmospheric composition during the first billion years of our planet’s history are just as debated as interpretations from the earliest records from the rocks, but a primordial history dominated by a CO_2_ and N_2_-rich secondary degassed atmosphere is plausible (e.g **Gaillard and Scaillet, 2014; Catling and Kasting, 2017**). Planetary dynamic models suggest that at times the early Hadean atmosphere was affected by temporary H₂-rich reducing conditions in the wake of large impacts. Metallic iron from asteroids and planetesimals reacting with surface water or oxidized mantle can generate transient H_2_-rich conditions that would be conducive for prebiotic synthesis of HCN and other reduced C-N species, before the impact flux waned (**Genda et al., 2017; Benner et al., 2020**). Photochemical models indicate that HCN production scales linearly with methane concentration, with [HCN] = 0.13 ± 0.01 × [CH₄] in upper-atmosphere analogs, though surface concentrations are expected 2–3 orders of magnitude lower (**Pearce et al., 2022**). For example, reduced atmospheres (from metallic asteroids, planetesimals or planetary embryos) complemented by carbonaceous chondrite-type materials can supply HCN at rates sufficient to produce nucleobases in ponds at concentrations of ∼1– 10 mM under favorable conditions (**Pearce et al., 2017**).

### 2.2 The Upper Kinetic Bookend – Isotopic and Fossil Evidence

The timing of life’s emergence is directly constrained by the earliest biosignatures, but interpretation of these signals remains contentious. For example, signatures of life are documented in ≥3.83 Ga paragneiss in Akilia association and ca. 3.8 Ga rocks of the Isua supracrustal belt, both located in the Itsaq Gneiss Complex of southern West Greenland (**Schidlowski, 1988; Mojzsis et al., 1996; Rosing, 1999; Manning et al., 2006; Schopf et al., 2007**). In the rocks of sedimentary origin reported by **Mojzsis et al. (1996)**, it was reported that carbonaceous inclusions within phosphate (apatite) minerals exhibit δ¹³C values of approximately -20 to -35‰ relative to PDB standard. The observed carbon isotopic fractionation is consistent with biological carbon fixation, as no known abiotic process can produce such light carbon and moreover, selectively associate the carbon with phosphate (**Mojzsis, 2026**). However, subsequent reappraisals stridently challenged these results and questioned whether some samples – especially from Akilia Island – may instead be 150–200 Myr younger, that metamorphism complicates the interpretation of traces of life, the quartz-pyroxene-magnetite rocks are instead metasomatized products of originally ultramafic lithologies, and/or that there may not even be graphite in these rocks (**Moorbath, 1998; Fedo et al., 2006**). A review of the controversies and the resolution of the conflict surrounding the interpretation of the Akilia rocks is provided elsewhere (**Mojzsis, 2026**). More controversial still is a 4.10 ± 0.01 Ga zircon from Jack Hills containing graphite with δ¹³C = −24 ± 5‰, interpreted as potentially biogenic (**Bell et al., 2017**). If it is evidence for life, this datum would confirm the presence of a metabolically sophisticated biosphere well into the Hadean. The crux of the debate over such lines of evidence stems from uncertainties in distinguishing biogenic from abiogenic carbon isotope fractionation in ancient, metamorphosed samples. Microfossil evidence captured in less altered sedimentary rocks provides less ambiguous albeit younger constraints. Structurally complex microfossils dated to ∼3.4–3.5 Ga (**Forterre, 2015; Schopf, 2006; Kaźmierczak and Kremer 2019)** seem to show that sophisticated microbial communities existed. The complexity of these microfossil structures points to organisms with a substantial prior evolutionary history. The oldest macrofossil shapes, in the form of bio-sedimentary structures termed stromatolites, are as old as the oldest microfossil evidence (e.g. **Walter et al., 1980**), whereas claims of yet older stromatolites from ca. 3.71 Ga rocks of the Isua Supracrustal Belt are confounded by the structural history of the rocks in question and mimicry by competent folds and ersatz stromatolite-like structures (**Zawaski et al., 2020**). Thus, by at least 700 Myr after Solar System formation we have in our archives tentative isotopic hints at a biosphere and by about 1 Gyr, the confirmed status of a thriving microbial community – plausibly including photosynthesizers – covering the Earth’s surface.

### 2.3 Quantitative Constraints on the Abiogenesis Timeframe

As shown above, the interval between the onset of continuous biocompatible conditions and the earliest biosignatures spans 130-900 Myr depending on which boundaries are accepted (**Figure 1**). The minimum interval (∼130 Myr) holds that stable conditions for prebiotic chemistry began at 4.33 Ga and modern life in the Domain Bacteria existed by ∼4.2 Ga (LUCA’s age from molecular clocks). The maximum interval (∼900 Myr) assumes conditions suitable for life almost immediately followed post-Moon-formation and that robust and interpretable microfossil/isotopic evidence for biological activity only exists after about 3.6 Ga. Bayesian statistical analyses of the oldest biosignatures provide moderate support (odds factors of a few to >10×, depending on which early signals are accepted) that abiogenesis is rapid owing to life’s early appearance (**Pearce et al., 2018; Spiegel and Turner, 2012**). These inferences are sensitive, however, to assumptions about the prior probability distribution for abiogenesis timescales and the reliability of Hadean biosignature claims. Some of these parameters are subjective and are as contingent and mutable as the available data.

**FIGURE 1:**
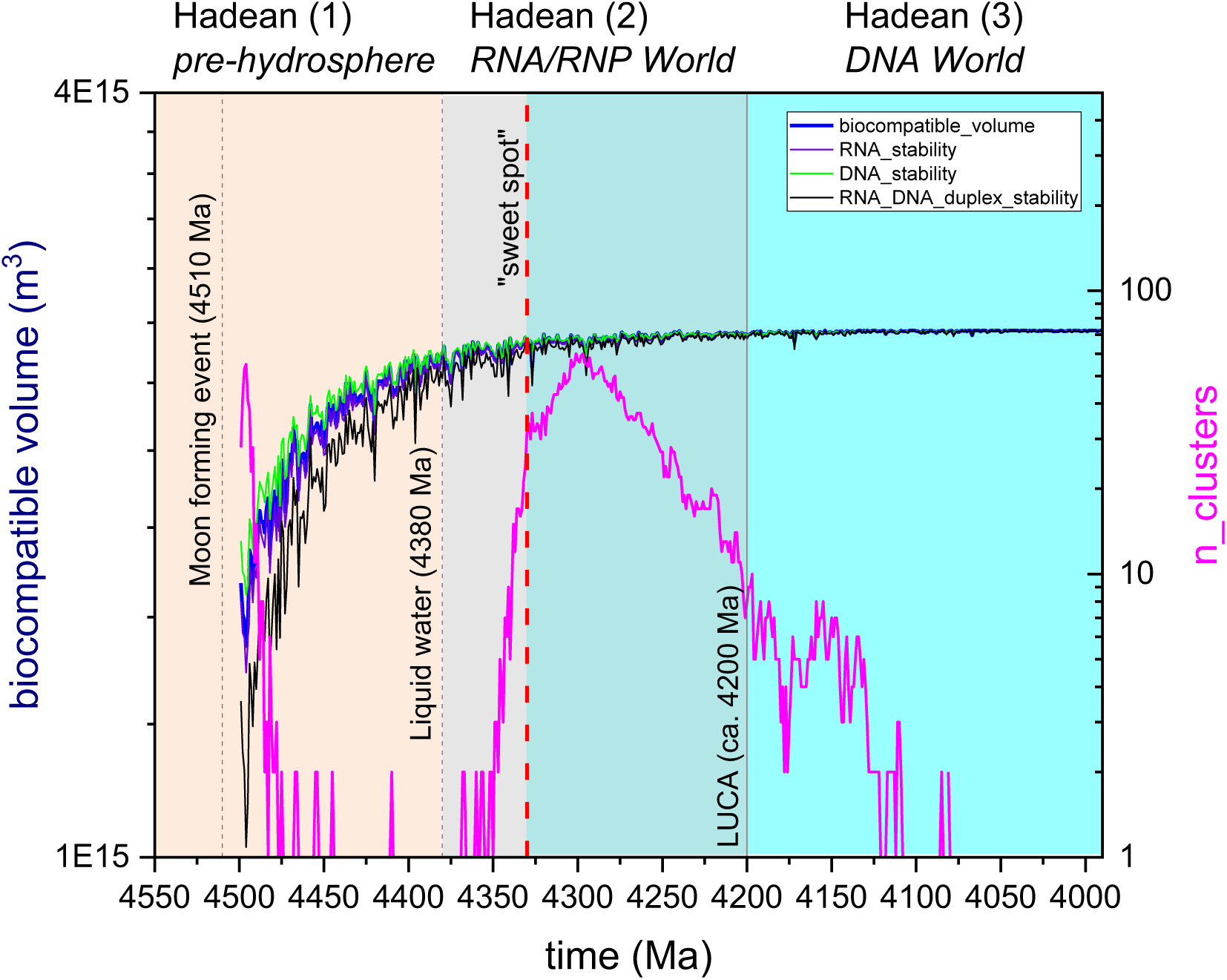
Hadean Timeline of Abiogenesis Windows. Geological timeline showing key events and kinetic windows for abiogenesis on Earth against biocompatible volumes (in m^3^) computed from thermal analysis of late accretion to the crust and the number of interconnected impact-generated hydrothermal systems (**Abramov et al., 2026**). Horizontal axis spans 4550-4000 Ma. Key events marked: **Hadean (1)** Moon-forming impact (4510 Ma; **Barboni et al. 2017**), **Hadean (2)** “sweet spot” or onset of continuous biocompatibility (4330 Ma), and **Hadean (3)** conforming to estimates of LUCA’s age (4200 Ma, 4.09–4.33 Ga range shown as shaded band; **Moody et al., 2024**). Colored bands indicate kinetic windows: pre-hydrosphere surface (4500-4400, orange), window for appearance of liquid water (4400 Ma onwards, light grey), and DNA-based cellular life (4.2 Ga onward, cyan). Overlapping bands illustrate temporal coordination of transitions.

The molecular clock constraint placing LUCA at ca. 4.2 Ga is particularly significant, as it implies that not only was modern-type DNA life present, but it had evolved to Bacteria-grade complexity with a ∼2.5 Mb genome encoding ∼2,600 proteins (**Moody et al., 2024**). Yet, we must also view this estimated date for LUCA with caution. Putting dates at the very root of the phylogenetic tree of all life relies heavily on mathematical assumptions. One of the problems encountered in the estimate of LUCA is that errors automatically propagate and multiply as you work backward toward the root node. Even with these caveats, the origin of life seems to occur once favorable conditions were established, and hence we argue that the kinetic barriers to cellular life appear to be surmountable on timescales of 10⁸ years or less.

### 2.4 Extrinsic Biocompatibility: Impact-Driven Chokepoints and Episodic Sterilization

The “Moneta hypothesis” proposes a ∼10²³ kg late veneer impactor (**Genda et al., 2017**) as the last major planet-wide sterilizing event after Moon formation, with peak plausibility for RNA synthesis occurring ∼120 ± 100 Myr after this impact, corresponding to ∼4.36 ± 0.10 Ga (**Benner et al., 2020**). Recent analysis has narrowed the “sweet spot” for abiogenesis to 4.33 Ga (**Abramov et al., 2026**), While model driven, these scenarios illustrate how impact-driven bottlenecks could episodically reset prebiotic progress, affecting when continuous habitability and uninterrupted evolution could begin (**Zahnle and Sleep, 2006**). It is worth mentioning that double-impact scenarios requiring paired impacts to deliver complementary reagents (**Anslow et al., 2025**) would be possible only very early in the history of the Solar System when multiple impacts were still common (>4.4 Ga; **Abramov et al., 2026**). This constraint suggests that abiogenesis after 4.4 Ga relied mostly on sustained geochemical sources of precursors rather than stochastic impact delivery.

While the threat of global sterilization over by ∼4.45 Ga, regional thermal stresses from large impacts associated with late accretion continued to affect the surface throughout the Hadean and early Archean. If early life involved an RNA biome inhabiting mild aqueous near-surface regions, such fragile proto-organisms would be susceptible to extinction from regional thermal perturbations. This scenario suggests that thermally robust DNA-peptide microbial communities adapted to deep crustal hydrothermal settings may have been better positioned to survive waning bombardment stages and repopulate the planet from deep subsurface refugia (e.g. **Abramov et al., 2013**).

## 3. Defining Kinetic Windows

### 3.1 Conceptual Foundation

We specify a kinetic window as being the temporal interval during which a specific set of physical-chemical conditions permits a particular reaction or process to proceed at rates sufficient to achieve biologically relevant outcomes. Unlike thermodynamic favorability (ΔG < 0), which determines whether a reaction can occur at equilibrium, kinetic accessibility (determined by activation energy *E_a_* and reaction rate (*k*) determines whether it will occur within the available time.

For abiogenesis, we postulate that multiple kinetic windows must be present on or in a planet’s crust, but not necessarily at the same time and location.

1. **Precursor synthesis**: Conditions permitting formation of nucleotides, amino acids, and lipid precursors at concentrations sufficient for subsequent assembly (typically >10 µM to 1 mM) (**Saladino et al., 2015**). In this framework, individual kinetic windows correspond directly to the environmental conditions required for the emergence of Gánti’s **(1997)** three essential subsystems: precursor synthesis and wet-dry cycling enable the genetic template, cellular organization supplies the boundary membrane, and functional metabolic emergence drives continuous energy flux. Probably only when all these distinct kinetic windows temporally and spatially overlap could the prebiotic chemical ensemble cross the critical threshold into a self-sustaining system capable of Darwinian evolution.
2. **Polymerization**: Conditions enabling oligomerization of monomers into functional polymers (RNA, peptides) without excessive degradation. For RNA, this requires balancing synthesis rates against hydrolysis rates, which depend exponentially on temperature and pH (**Li and Cantor, 1971; Levy and Miller, 1998**).
3. **Functional emergence**: Conditions allowing evolution of catalytic activity (ribozymes, peptides) and self-replication. This requires sufficient molecular diversity (sequence space exploration) and stability (preservation of functional sequences) (**Lehmann and Joyce, 1993**).
4. **Cellular organization**: Conditions permitting encapsulation, compartmentalization, and the emergence of protocells with heredity and metabolism (**Szostak et al., 2001; Luisi et al. 2006**).
5. **Evolutionary innovation**: Conditions supporting the transition from RNA to ribonucleoprotein (RNP) to DNA-based genomes, requiring sustained environments for selection and fixation of innovations (**Lazcano and Miller, 1996**).

Each kinetic window is characterized by: - **Duration (Δt)**: The time interval during which conditions remain favorable, ranging from 10³ years (pond evaporation cycles) to 10⁸ years (geological epochs). – **Spatial extent (V)**: The volume or area over which favorable conditions exist, from microscopic pores to planetary-scale environments. – **Parameter ranges**: The specific values of temperature (K), pH, water activity (*a_w_*), UV flux (Φ), and elemental concentrations ([X]) that define favorability. – **Recurrence frequency (*f*)**: For cyclic processes (wet-dry cycles, diurnal/seasonal variations), the frequency at which favorable conditions repeat. We return to these characteristics in Sec. 8 with worked examples for different planetary environments.

### 3.2 Rate-Limiting Steps and Bottlenecks

The overall rate of abiogenesis is determined by the slowest (rate-limiting) step in the sequence. Identifying these bottlenecks is necessary for understanding which environmental parameters exert the strongest control on timescales. Experimental and modeling studies have identified several candidate rate-limiting steps. Ribose is thermodynamically unstable with half-lives of days to weeks at moderate temperatures (25–50°C) (**Larralde et al., 1995)**. Achieving concentrations >10 µM required for nucleotide assembly necessitates either rapid synthesis or stabilization mechanisms such as by borate complexation or mineral adsorption (**Breslow, 1959; Ricardo et al., 2004**). Condensation of nucleobases, ribose, and phosphate to form nucleotides is thermodynamically unfavorable in water (ΔG > 0) and requires the require activation chemistry or dehydration (**Orgel, 2004**). Wet-dry cycling in ponds can drive this reaction but necessitates specific pH (8–10) and mineral catalysts (**Ferris et al., 1996; Powner et al., 2009**). Template-directed RNA synthesis by ribozymes exhibits error rates of ∼10⁻² to 10⁻³ per nucleotide (**Bartel and Szostak, 1993**), which limits the maximum length of accurately replicated sequences to ∼100–1000 nucleotides without error correction. This “error catastrophe” threshold constrains the complexity achievable in a pure RNA World (**Eigen, 1971**). The transition from RNA to RNP systems requires coevolution of RNA catalysis with peptide synthesis, or perhaps there was no RNA World and only a RNP World existed as the precursor form (**Müller et al., 2022**). Short cationic peptides can stabilize RNA structures and enhance ribozyme catalytic rates by 10–100-fold (**Tagami et al., 2017**), creating a positive feedback loop, but the initial emergence of this synergy is kinetically challenging (**Vaidya et al., 2012**). Spontaneous vesicle formation from amphiphilic molecules requires concentrations >10⁻⁴ M and specific lipid compositions (**Walde et al., 1994**). Maintaining membrane integrity during growth and division while allowing nutrient influx is a kinetic balancing act (**Chen et al., 2004; Walton et al., 2026**). It may be possible to overcome this last inhibiting step by concentrating and stabilizing abundant and diverse protocell structures on mineral surfaces (**Köksal et al, 2022**).

### 3.3 Environmental Sensitivity and Parameter Coupling

Kinetic windows are highly sensitive to environmental parameters, often exhibiting threshold behavior where small changes in conditions cause large changes in rates. These parameters are often coupled: changing temperature affects pH, water activity and reaction rates. Reaction rates typically follow Arrhenius behavior, k = A exp(−Ea/RT), doubling every ∼10°C for typical activation energies (Ea ∼ 50–100 kJ/mol). Degradation rates also increase with temperature, creating an optimal temperature range for net synthesis. Also, RNA hydrolysis rates increase exponentially with pH above 7 due to hydroxide-catalyzed cleavage of the phosphodiester backbone (**Li and Cantor, 1971**). Below pH 5, acid-catalyzed depurination becomes significant. This creates a narrow pH window (6–8) for RNA stability (**Figure 2**). Biochemical reactions require a*_w_* > 0.6 for enzyme activity (**Grogan and Zachara, 1999**), but polymerization reactions are thermodynamically favored at low a*_w_* (dehydration). Wet-dry cycling resolves this paradox by alternating between synthesis (dry) and function (wet) phases (**Deamer and Barchfeld, 1982**). Lastly, UV radiation (200–300 nm) damages nucleic acids through pyrimidine dimer formation and strand breaks, with damage rates proportional to flux (**Cadet et al., 1990**). However, moderate UV may drive prebiotic synthesis of some precursors (**Todd et al., 2021; cf. Miller, 1953**). The optimal UV flux depends on the balance between synthesis and damage, which remains to be better quantified. Indeed, this is an old problem. Many are unaware that Stanley Miller actively considered UV radiation in prebiotic chemistry. While he and Harold Urey decided to use electric discharges for their iconic 1953 experiments owing to the relative experimental ease, they both understood that the UV flux from the young Sun was also an important and relevant energy source for prebiotic chemistry.

**FIGURE 2.**
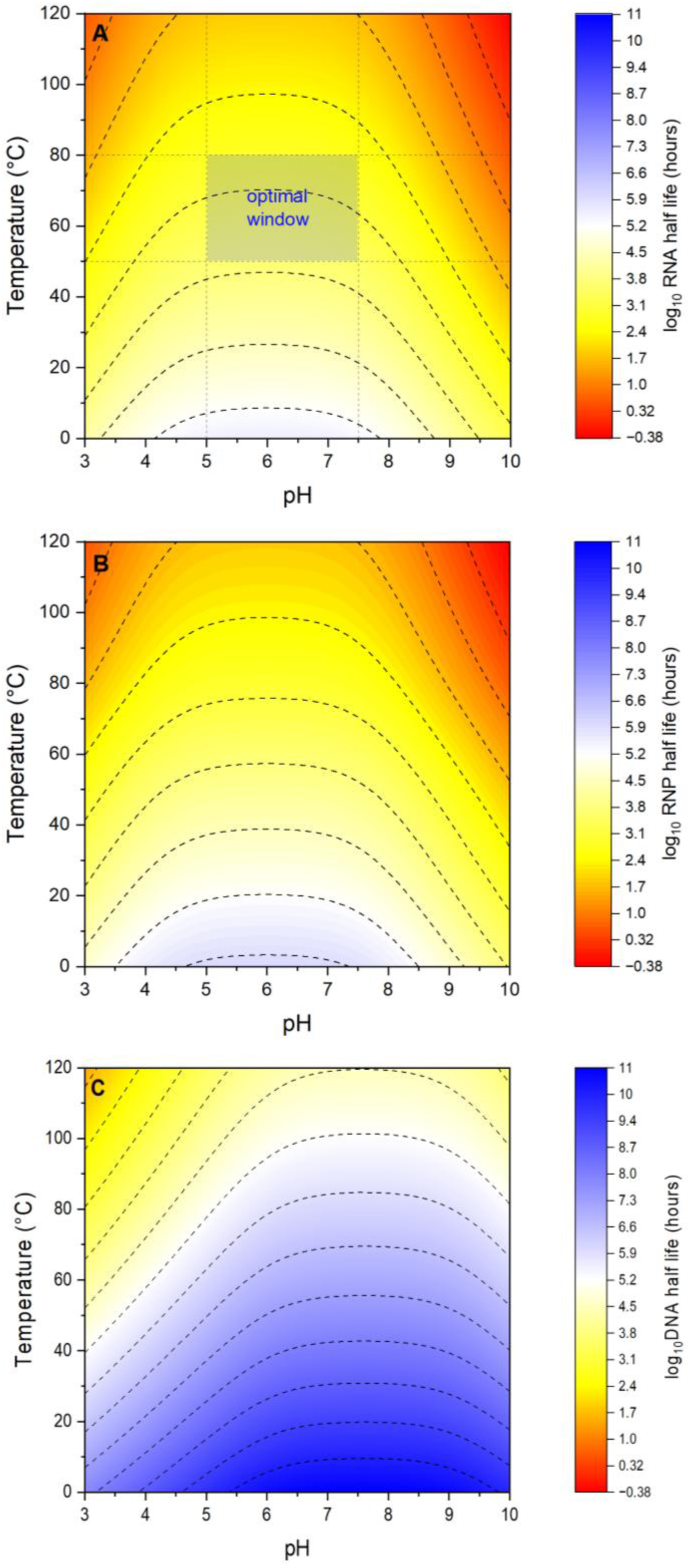
Physical-Chemical Parameter Spaces for RNA, RNP and DNA Stability on Hadean Earth. Three-dimensional parameter space showing kinetic windows for RNA (**A**), RNP (**B**) and DNA (**C**) stability as a function of temperature (0–120°C) and pH (3–10). Color gradient indicates half-lives (log_10_ scale) in hours: blue (>10 years), yellow (>10 days), orange (<day), red (minutes to hours). Optimal window for RNA is relatively narrow: T = 50–80°C, pH = 6-7.5, and [Mg²⁺] = 1–10 mM.

### 3.4 Temporal Coordination and Sequential Windows

Different stages of abiogenesis that satisfy Gánti’s chemoton require different environmental conditions, implying that kinetic windows must occur in sequence rather than simultaneously. For example, nucleotide synthesis is favored at pH 8–10 and moderate temperatures (50–80°C) with wet-dry cycling (**Powner et al., 2009**). Also, RNA stability is maximized at pH 6–7 and lower temperatures (0–30°C) (**Hud et al., 2013**). Ribozyme catalysis requires Mg²⁺ concentrations of 1–10 mM (**Cech, 2000**), but Mg²⁺ also catalyzes RNA degradation (**Brion et al., 1999**). Peptide synthesis on ribosomes requires temperatures <100°C and near-neutral pH (**Noller, 2005**). Finally, DNA synthesis and replication are favored at moderate temperatures (30–60°C) and pH 7–8 (**Kornberg and Baker, 1992**). This sequential requirement implies that abiogenesis likely involved migration or fluid mixing between environments (e.g., from shallow alkaline hydrothermal vents for synthesis to cooler, near-to super-neutral ponds for stability and function) or temporal evolution of a single environment (e.g., cooling and pH neutralization of early Earth’s surface) (**Russell, 1997**). The duration of the abiogenesis window is thus determined not only by the persistence of any single set of conditions but by the coordination of the entire sequence in time.

## 4. Physical-Chemical Parameter Spaces for Biocompatibility

### 4.1 Temperature: Stability versus Kinetics

Temperature exerts dual control on abiogenesis: it accelerates both synthesis and degradation reactions, creating an optimal range that balances productivity against stability. For RNA-based systems, this trade-off is especially acute compared to RNP- or DNA-based life. The phosphodiester backbone of RNA undergoes base-catalyzed hydrolysis with a rate constant that increases exponentially with temperature. For example, at pH 7 and 25°C, the half-life (τ_½_) of RNA is 10^4.5^ hours or ∼3.6 years, but this decreases to ∼6 months at 50°C and ∼1 month at 75°C (**Levy and Miller, 1998**). Above 100°C, RNA is unstable on timescales of days to weeks, precluding the existence of a high-temperature RNA World (**Joyce, 2002**). Experimental studies of ribozyme activity and stability suggest an optimal temperature range of 0–50°C for RNA-based life (**Matthews et al., 1999)**. This is consistent with reconstructions of LUCA as mesophilic or moderately thermophilic (not hyperthermophiles), based on the absence of reverse gyrase – an enzyme systematically present in all hyperthermophiles – in the universal protein set (**Forterre, 2002**). Reconstructed rRNA and protein sequences also support a mesophilic LUCA (**Lane et al., 2010; Galtier et al., 1999**). Since RNA is inherently more chemically unstable than DNA at extreme temperatures, it has long been a scientific consensus that RNA viruses struggle to survive in hyperthermophilic hosts (like Archaea, which thrive at temperatures above 80°C). All currently isolated hyperthermophilic viruses rely on highly stable, double-stranded DNA genomes to survive the intense heat of their hosts’ native environments. Metagenomic studies of high-temperature acidic hot springs, however, have detected distinct RNA viral genome fragments. That said, no hyperthermophilic RNA virus has ever been successfully isolated or cultured to date (**e.g. Bolduc et al., 2012**). This leaves the question open as to whether RNA viruses are present in high temperature geologic settings that possibly inhabit hosts we cannot cultivate, or that RNA cannot survive these conditions.

Geochemical models, climate simulations and zircon oxygen isotope data suggest that Hadean surface temperatures may have been elevated relative to modern values but not uniformly extreme. Early estimates of >200°C surface temperatures (**Sleep et al., 1989**) have been revised downward based on evidence for liquid water and continental crust by ∼4.4 Ga (**Mojzsis, 2026**). A plausible scenario involves surface temperatures of 50–100°C in the early Hadean (4.45–4.35 Ga), declining to near-modern values (0–30°C) by the Hadean-Eoarchean transition at 3.9–3.8 Ga (**Zahnle et al., 2010**). This hypothetical cooling trajectory has implications for the RNA World. Initially high temperatures (>80°C) would preclude stable RNA, limiting prebiotic chemistry only to more thermally robust precursors (amino acids, simple organics). As temperatures declined into the 30-80°C range, an RNA World became kinetically feasible (**Abramov et al., 2026**).

Submarine hydrothermal vents provide localized high-temperature environments (100–400°C) that persist throughout Earth’s history wherever hot rock meets water (**Martin et al. 2008**). While too hot for RNA stability, these systems offer chemical disequilibria (H_2_, reduced metals, pH gradients) that can provide the first steps towards autotrophy (**Russell and Hall, 1997; Lane and Martin, 2012**). Alkaline hydrothermal vents (pH 9-11, T ∼ 40-90°C) represent a particularly compelling environment at the surface or in the shallow subsurface, combining moderate temperatures with conditions conducive to prebiotic chemistry and nascent autotrophy (**Kelley et al., 2005; Russell et al., 2014**). But, as shown here, these pH and temperature conditions are sub-optimal for purely RNA- or even RNP-based life.

Further afield, carbonaceous asteroids experienced transient heating from ²⁶Al decay, with habitable temperatures (273–373 K) maintained for ∼24 Myr in the center of a 200-km body with 40% initial ice content (**Abramov and Mojzsis, 2016**). Yet, alteration temperatures in CM meteorites (273–308 K) are at the lower end of the range favorable for prebiotic chemistry, and the brief duration of warm conditions (∼10⁷ years) limits the extent of chemical evolution possible.

### 4.2 pH: The Alkaline Hydrolysis Problem

After temperature, the most critical parameter for RNA stability is pH because hydroxide ions catalyze cleavage of the phosphodiester backbone through nucleophilic attack on the phosphorus atom facilitated by the adjacent 2’-hydroxyl group (**Saenger, 1984**). As previously mentioned, the rate of alkaline hydrolysis increases by approximately one order of magnitude per pH unit above 7. At 25°C, RNA half-life is 10^4.2^ hours (∼1.7 years) at pH 8, 2.5 months at pH 9, and 8 days at pH 10 (**Oivanen et al., 1998**). This exponential dependence creates a narrow pH window (6–8) for RNA stability at low temperatures on timescales relevant to chemical evolution (decades). Below pH 5, acid-catalyzed depurination (loss of purine bases) becomes significant (**Lindahl and Nyberg, 1972**), further constraining the viable range for RNA to persist in a prebiotic (or not-completely-alive, *para*biotic) setting. Early Earth’s ocean pH is estimated to have been acidic (∼5.0) due to high atmospheric CO₂ levels and limited carbonate buffering (e.g. **Krissansen-Totten et al., 2018**). At 10-100× present atmospheric CO_2_ level, the equivalent partial pressure of CO_2_ would be 0.004-0.04 bar, respectively; compare this to Venus with 89 bars of CO_2_. As surface temperatures declined and silicate weathering accelerated towards the end of the Hadean and into the Eoarchean, CO_2_ was drawn down and ocean pH rose toward neutral values (**Walker, 1985**). This pH rise had paradoxical effects on the RNA World. Initially acidic conditions (pH 5-6) were suboptimal for nucleotide synthesis but provided moderate RNA stability. As pH rose toward circum-neutral (pH 7-8) or higher, nucleotide synthesis became more favorable, but RNA half-lives shortened. Further pH rise above 8 progressively destabilized exposed ribozymes, which we speculate was an environmental selection factor against the RNA World.

Furthermore, experimental studies indicate that nucleotide phosphorylation and polymerization are favored at mildly alkaline pH (8–10) in the presence of mineral catalysts (**Ferris, 2006**). Alkaline hydrothermal systems (pH 9–11) thus emerge as plausible sites for nucleotide synthesis, despite being suboptimal for RNA stability. This evokes some kind of spatial or temporal separation between synthesis (alkaline environments) and function (neutral environments) such as might be expected in mixing pools and drainages from different surface environments in close geographic proximity (**Powner and Sutherland, 2011**). Divalent magnesium is essential for RNA catalysis, stabilizing tertiary structures and facilitating phosphoryl transfer reactions (**Piccirilli et al., 1993**). However, Mg²⁺ also catalyzes RNA degradation, particularly at alkaline pH where Mg(OH)⁺ acts as a nucleophile (**Brion et al., 1999**). The optimal Mg²⁺ concentration depends on pH: at pH 7, 1–10 mM Mg²⁺ is beneficial, but at pH 9, even 1 mM Mg²⁺ significantly accelerates degradation. This coupling between pH and metal ion chemistry further constrains the viable parameter space for the RNA World. Arguably, marine environments (pH ∼8, high Na⁺ and Mg²⁺) are unsatisfactory for RNA stability due to alkaline hydrolysis and metal-catalyzed degradation (**Szostak, 2012**). As is evident here, we favor freshwater ponds on emergent land, with lower pH (6–7) and reduced metal concentrations, to provide more favorable conditions (**Pearce et al., 2017; Damer and Deamer, 2020; Sahai et al., 2022**). The presence of exposed landmasses and meteoric water-crust interactions in the Hadean eon thus created prime habitats for the RNA and RNP Worlds.

It is noteworthy that alkaline crater lakes and evaporative soda basins (pH 8–10) provide favorable environments for early prebiotic chemistry. These settings are typically rich in sodium, dissolved carbonate and bicarbonate, and small amounts of borates, while containing low levels of divalent metal ions (**Kempe and Kaźmierczak, 1990**). Endorheic systems such as these can concentrate important prebiotic compounds (e.g. HCN derivatives, formamide, and cyanamide). They may also support borate-mediated stabilization of D-ribose, protecting it from thermal decomposition (**Ricardo et al 2004**).

### 4.3 UV Radiation: Damage versus Synthesis

If the Hadean atmosphere was transparent and not too dense, we can expect that ultraviolet radiation (200–300 nm) reaching the surface from the young Sun was significantly more intense than today, with fluxes estimated at 10-100× present levels (**Cockell, 2000**). Ultraviolet light exerts opposing effects on abiogenesis: it damages nucleic acids through pyrimidine dimer formation and strand breaks, but it also drives photochemical synthesis of some prebiotic precursors. We now examine some of the repercussions of an intense flux. Pyrimidine dimers form when adjacent pyrimidine bases (cytosine, uracil) absorb UV photons and undergo [2+2] cycloaddition (**Cadet, 1990**). These dimers block replication and transcription, with damage rates proportional to UV flux. At modern surface UV levels, unprotected RNA accumulates ∼1 dimer per 10⁴ nucleotides per hour (**Mitchell and Nairn, 1989**). At 10× modern flux, this increases to ∼1 dimer per 10³ nucleotides per hour, requiring repair mechanisms or some kind of shielding such as pigment-like molecules or minerals, for RNA survival. The presence of RNA photolyase activity in LUCA implies that early life was exposed to UV radiation and evolved repair mechanisms (**Forterre, 2005**). Photolyases themselves, however, are complex enzymes requiring cofactors (flavin, folate), which we take to mean that UV exposure was a selective pressure after cellular life emerged rather than something that life had to endure during the prebiotic phase (**Sancar, 2003**).

If we are correct, this last point suggests that the utilization of pigments and ultimately photosynthesis was a relatively late innovation compared to early forms of heterotrophy and/or chemoautotrophy. Early life still had to contend with the energetic nature of the young Sun if prebiotic chemistry occurred near the surface, and several mechanisms can be invoked that shielded prebiotic chemistry from UV damage. In a pure water column, UV-C (200–280 nm) is attenuated within the top 1-10 m and is much more attenuated in seawater and turbid waters owing to absorption and scattering (e.g. **Wozniak and Dera, 2007**). Importantly, even shallow subsurface marine environments such as those near surface hydrothermal vents and the shallow seafloor provide natural UV shielding. Adsorption of nucleic acids onto mineral surfaces (clays, zeolites) reduces UV damage by 10-100-fold because the photons are absorbed by the shielding mineral (**Biondi et al., 2017; Scappini et al., 2004)**. Photochemical hazes in a reducing atmosphere also absorb UV, reducing surface flux (**Trainer et al., 2006; Zahnle, 1986**).

One way of interpreting the presence of photolyases in the LUCA is that it is an ancient relict. Prebiotic chemistry in shallow ponds may have been able to exploit diurnal cycles with synthesis occurring at night and UV exposure during the day driving selection for UV-resistant sequences (e.g. **Pearce et al., 2017**). It was shown long ago that UV photolysis of simple molecules (CH_4_, NH_3_, H_3_O, CO_3_) generates reactive intermediates like radicals and excited states which drive synthesis of amino acids, nucleobases, and other organics (**Miller and Urey, 1959**) and highlighted more recently by **Chyba and Sagan (1992)**. The net effect of UV on abiogenesis thus depends on the balance between synthesis and damage, which varies with atmospheric composition, water depth, and the stage of chemical evolution.

### 4.4 Water Activity: The Dehydration Paradox

Water activity (*a_w_*) is the ratio of the vapor pressure of water in a solution to that of pure water under the same conditions, ranging from 0 (completely dry) to 1 (pure water). Biochemical reactions generally require *a*_w_ > 0.6 for enzyme activity and membrane integrity (e.g. **Grogan and Zachara, 1999**). Yet at the same time, polymerization reactions including peptide bond formation and phosphodiester bond formation are thermodynamically unfavorable in water due to the opposing hydrolysis reaction (**Orgel, 1998**). While there exists no debate whatsoever that life, including pre-life, requires water as a solvent, the synthesis of its key polymers (proteins, nucleic acids) is inhibited by water. This paradox is resolved through several mechanisms existing in the natural environment (e.g. **Mojzsis et al., 1999**):

1. **Wet-dry cycling**: Evaporation of ponds concentrates reactants and reduces *a*_w_, driving polymerization during the dry phase. Rehydration during the wet phase allows diffusion, mixing, and functional activity.
2. **Eutectic freezing**: Freezing of dilute solutions concentrates solutes in liquid veins between ice crystals, effectively reducing *a*_w_ and driving polymerization (e.g. **Damer and Deamer, 2015**). This mechanism is relevant for cold environments (polar regions, Hadean ice world scenarios, **Zahnle, 2006**).
3. **Mineral or glass surfaces**: Adsorption onto mineral surfaces can reduce the local water activity and orient monomers for polymerization (**Cairns-Smith, 1982**). Basaltic glass catalyzes RNA polymerization (**Jerome et al., 2022**) perhaps in part by providing a rich surface charge and excluding bulk water (**Ghiara et al., 1993**).

Here, we purposefully exclude using high-energy intermediates (e.g., nucleoside triphosphates, aminoacyl adenylates) to shift the thermodynamic equilibrium toward polymerization (e.g. **Schwartz and Orgel, 1985**). This synthesis requires a source of chemical energy (ATP, pyrophosphate) that must itself be synthesized.

Up to this point, we have emphasized the importance of exposed landmasses, rainfall, and evaporative ponds on Hadean Earth to host settings for wet-dry cycling. We have not mentioned previously how high salinity (>0.5 M NaCl) disrupts RNA secondary structures by screening electrostatic interactions (**Santa-Lucia Jr, 1998**). Dilute or brackish settings (0.01–0.1 M) are thus favored for the RNA World (**Szostak, 2012**). This constraint alone favors freshwater ponds over marine environments, consistent with the low-metal requirements (e.g. **Sahai et al., 2022**). Diurnal and seasonal variations in temperature and humidity would drive repeated cycles, with each cycle potentially adding more nucleotides to growing oligomers. Over 10⁶–10⁷ cycles (speculatively corresponding to 10³-10⁴ years), functional ribozymes could have emerged.

## 5. The RNA World: Duration, Constraints, and Kinetic Bottlenecks

In brief, the RNA World hypothesis postulates that early life consisted of RNA as both genetic material and catalyst, preceding the evolution of DNA and proteins (**Gilbert, 1986; Joyce, 2002**). This hypothesis is supported by the central role of RNA in modern cells (mRNA, tRNA, rRNA, ribozymes) and the discovery of catalytic RNAs capable of self-replication and metabolism (**Cech et al., 1981; Guerrier-Takada et al., 1983**). However, recent work suggests that the RNA World was a brief, transient stage rather than an extended epoch. Molecular clock analyses placing LUCA at ∼4.2 Ga combined with the onset of continuous habitability at ∼4.33 Ga, constraining the duration of the RNA and RNP Worlds to less than 120 Myr before the LUCA (**Abramov et al., 2026**). Within the framework of the kinetic windows hypothesis presented here, such a tight timeline is consistent with the intrinsic chemical instability of RNA and the kinetic advantages of transitioning to DNA-based genomes.

### 5.1 Chemical Instability of RNA

As reviewed elsewhere (**Medvegy et al. submitted**) RNA’s chemical instability primary arises from such detrimental shortcomings as sensitivity to alkaline hydrolysis, depurination and metal-catalyzed degradation. In alkaline hydrolysis, the 2’-hydroxyl group on ribose acts as an intramolecular nucleophile, attacking the adjacent phosphorus atom in the phosphodiester backbone. At acidic pH (<5), the N-glycosidic bond linking purine bases (adenine, guanine) to ribose undergoes acid-catalyzed hydrolysis, releasing the base and leaving an abasic site. Depurination rates increase with temperature, with half-lives (τ_½_) of <4 years at pH 5 and 25°C, decreasing to ∼6 months at 50°C. This mechanism limits RNA stability in acidic environments and at elevated temperatures compared to RNP (τ_½_ = ∼1.2y at 50°C and pH 5) or DNA (τ_½_ = ∼440y at 50°C and pH 5). Divalent metal ions, particularly Mg²⁺, are essential for RNA catalysis but also catalyze degradation. The Mg²⁺ coordinates with the phosphate backbone and facilitates nucleophilic attack by water or hydroxide, accelerating cleavage. At pH 7 and 1 mM Mg²⁺, this effect is modest, but at pH 9 and 10 mM Mg²⁺, RNA half-life decreases to weeks (**Brion et al., 1999**). It is interesting to note that Fe²⁺ can partially substitute for Mg²⁺ in catalysis (**Jaeger et al., 2006**) but also catalyzes degradation through Fenton chemistry with the generation of hydroxyl radicals (**Halliwell and Gutteridge, 1984**).

Certain hot springs hosted in ocean island basalt (pH ∼8; 55-75°C) mirror the characteristics cited above. One example of this is the hot spring in Hveragerdi, Iceland approximately 45 km east of Reykjavik investigated by **Sonne-Hansen and Ahring (1997)**. Saliant concentrations of the various ions in the pool water were as follows: Ca^2+^ 5 mg l^−1^, Mg^2+^ < 1 mg l^−1^, Na^+^ 23 mg l^−1^, K^+^ 2.3 mg l^−1^, SiO_2_ 116 mg l^−1^, Fe 0.42 mg l^−1^, Cl^−^ 23 mg l^−1^, PO ^3-^ 24 μg l^−1^, and HBO 17 μg l^−1^. The last constituent – metaboric acid – is noteworthy in this context. Borate stabilizes ribose sugars and facilitates the polymerization of amino acids into polypeptides under prebiotically plausible, neutral or acidic conditions (**Benner et al., 2010**; **Scorei, 2012**). Concentration of metaboric acid requires raising pH of the solution to >8 and evaporation to drive the equilibrium toward stable polyborate species (**Tsuyumoto et al., 2007**). We find this to be an interesting contemporary analogue chemistry. Chemical parameters seem to point to Mg^2+^ at low concentrations and Fe^2+^ at even lower concentrations at pH >8 and at mild temperatures under conditions allowing for evaporation to define the aqueous environment for the emergence of the RNA World.

### 5.2 Error Catastrophe and Genome Size Limits

The nature of RNA replication by ribozymes is error-prone, with per-nucleotide error rates of ∼10⁻² to 10⁻³ (**Bartel and Szostak, 1993**). This error rate imposes a limit on the maximum genome size that can be stably maintained, known as the Error Catastrophe threshold (**Eigen, 1971**).

Eigen’s error threshold can be viewed in the following way. For a genome of length L and per-nucleotide error rate ε, the fraction of error-free replicates is:

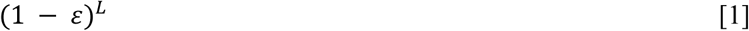

Stable inheritance requires:

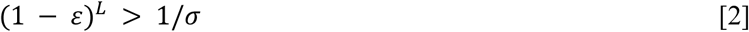

where *σ* is the selective advantage of the functional sequence (**Eigen and Schuster, 1977**). For *ε* = 10⁻², this limits *L* to ∼100 nucleotides; for *ε* = 10⁻³, *L* ∼ 1000 nucleotides (**Briebricher and Eigen, 2005**). A genome of 100–1000 nucleotides can encode only a few small ribozymes (typical ribozymes are 50–200 nucleotides; **Ferré-D’Amaré and Doudna, 1999**). This has interesting implications for RNA World complexity because it severely limits the metabolic and regulatory complexity achievable in a pure RNA World.

Indeed, according to **Moody et al. (2024)**, LUCA’s genome is at least three orders of magnitude larger than that of a hypothetical RNA organism which implies that the transition to DNA with its lower error rate due to proofreading and repair was required for any kind of meaningful genomic expansion (e.g. **Kunkel and Bebenek, 2000**). One proposed solution to the error catastrophe is organization of ribozymes into hypercycles, networks of mutually catalytic molecules that collectively maintain information (**Eigen and Schuster, 1979; Maynard Smith and Szathmáry, 1995**). Hypercycles are, however, vulnerable to parasites (i.e. non-functional sequences that replicate faster) and require spatial compartmentalization to be stable (**Szathmáry and Demeter, 1987; Boerlijst and Hogeweg, 1991**). Compartmentalization or encapsulation is a management strategy which effectively and controllably isolates the different components or groups of different components mitigating out-of-control complexity (**Mojzsis 2026**). In our view, a cell encapsulating the information apparatus is something that existed with the origin of the RNA World.

### 5.3 Proto-metabolism and Pseudo-Heterotrophy

We do not know how a very primitive pure RNA World organism could be powered by some kind of metabolism that directly benefited it in growth and function. In principle, the first steps to metabolism can be viewed quite classically by drawing on prebiotic organic pools as a food source (**Horowitz, 1945**) feeding into ribozymes, which catalyzed essential reactions for self-maintenance and replication (**Joyce 2002; Robertson and Joyce 2012**). Elsewhere, we have argued that early RNA life made direct use of abiotic exogenous synthesized in the environment which could pass through protocell membranes and directly interacted with simple RNA strands contained therein (**Medvegy et al. submitted**). Alternatively, direct absorption of formamide, cyanamide, or glycolaldehyde with ribozymes facilitating their condensation into nucleotides or energy-rich intermediates has also been proposed (**Damer and Deamer 2020**).

### 5.4 The Ribonucleoprotein (RNP) World was a Precursor to LUCA

The severe limitations inherent to a pure RNA World drove the evolution of peptide synthesis where RNA and peptides coevolved on the road to what is termed the ribonucleoprotein (i.e. RNP) world (**White, 1976**). The advantages gained by an RNP organism over pure RNA life is that short cationic and hydrophobic peptides can stabilize RNA secondary structures, enhance ribozyme catalytic rates by 10–100-fold, and provide new catalytic functionalities (**Tagami et al., 2017**). These synergistic advantages between peptide and RNA motivated the origin of the RNP World. This is because positively charged peptides (rich in lysine, arginine) bind to the negatively charged backbone of RNA, neutralizing electrostatic repulsion and stabilizing tertiary structures (**Erat and Sigel, 2011**). Hydrophobic peptides can insert into membranes, providing transport functions inaccessible to RNA (**von Heijne, 2006**) creating a positive feedback loop: RNA-encoded peptides enhance RNA catalysis, enabling more efficient peptide synthesis (**Vaidya et al., 2012**).

We envision that the first steps to the RNP World began with small “wild” peptides that formed abiotically in the environment which were able to pass through proto-cellular structures already containing encapsulated RNAs and find binding sites therein (e.g. **Medvegy et al. submitted; Gözen, 2026**). This is a variant of the Retrograde Hypothesis of **Horowitz (1945)**. While peptides can diffuse through protocell membranes, the ability to do so depends on size, charge, and sequence of the molecule. Passive diffusion of wild peptides ought therefore to have been highly selective into protocells. To get larger or highly charged peptides across membranes without active transport mechanisms unavailable in early protocells, these molecules relied on several specific avenues of entry. One of these is pore formation, where certain peptides self-assemble into higher-order structures that insert themselves into the membrane to create transient aqueous pores (or “toroidal pores”) that allow molecules to pass. Some amphipathic peptides are naturally capable of penetrating lipid bilayers directly or by causing temporary, localized membrane deformation. Some peptides can also bind directly to the membrane interface, altering its structural properties and making it more porous to other cargo (**Cooper and Hausman, 2000**). The idea here is that successful binding sites of wild peptides to early RNA or RNA-like structures yielded more stable systems which went on to survive long enough to effectively replicate and outcompete pure RNA organisms almost immediately. There may have been no need for a pure RNA World if life began with the structurally more stable RNP World (**Müller et al., 2022**).

The central molecular machine of the RNP World is the ribozyme (**Cech, 2000**). This molecular entity underwent an unimaginably complex evolutionary trajectory from primitive peptidyl transferase center (PTC)/tRNA systems to the multi-subunit structure in modern cells (**Fox, 2010; Petrov et al., 2015**). Phylogenetic analysis suggests that small subunit components (mRNA binding, translational processivity) evolved before large subunit catalytic centers (**Williams and Fecko, 2012**). An accretion model, where components were added incrementally with each conferring selective advantages, is consistent with the observed structure (**Petrov et al., 2014**).

Indications are that the LUCA possessed a ribosome with ∼30-40 proteins, equivalent to about half of modern ribosomes, but with 90% of the rRNA structure conserved between Archaea and Bacteria already having been set up beforehand (**Forterre, 2015**). The length of time available before the LUCA wherein a ribosome at this level of complexity was built is currently the subject of intense interest. The LUCA had 44-45 tRNAs with 2-3 modifications, reading 59-60 of 61 sense codons, and used the modern optimized genetic code. Yet, we now understand that translation was already accurate, meaning that it was capable of synthesizing bona fide proteins in contradiction to the “progenote” hypothesis of a primitive, error-prone LUCA (**Woese and Fox, 1977**).

## 6. The World of LUCA

It is worth noting that in our discussions of the nature of the LUCA, we attempt to avoid any description of it as being “prokaryotic”. The concept of the prokaryote was refuted in 1977 when Carl Woese discovered via comparisons of ribosomal RNA that a separate domain of life – Archaea – existed; he went on to propose a phylogenetic tree that related the most diverse forms of life, Bacteria and Archaea, to everyone else (**Pace, 1997**).

### 6.1 Cellular Apparatus

Analysis has shown that the LUCA possessed the core cellular apparatus of microbes of the contemporary Bacteria and Archaea domains (**Woese, 1998**). It likely had basic phospholipid membrane constituents, though the exact composition remains uncertain; it possessed genes for peptidoglycan synthesis (mraY, murC), indicating a cell wall (e.g. **van Heijenoort, 2001**). The cytoskeletal actin-like protein MreB was also present to provide structural support (e.g. **Jones, 2001**). It had membrane-bound ATP synthase subunits, including V/A-type subunits A (reported posterior probability PP = 0.98) and B (PP = 0.94), V/AO subunits I/a (PP = 0.99) and K/c (PP = 0.82), and F1-type β-subunit (PP = 0.94) (e.g. **Müller and M. Grüber, 2003**). This is consistent with many previous studies mapping ATP synthase to LUCA (**Hillis and Bull, 1993; Forterre and Philippe, 1999**) and indicates that it could generate ATP via chemiosmotic coupling. Translation was already accurate and capable of synthesizing bona fide proteins (**Forterre, 2015**). An early CRISPR-Cas system was in place with support for 19 class 1 effector protein families, including types I and III (**Moody et al., 2024; cf. Koonin and Makarova, 2019; Koonin et al., 2015**). For instance, Cas3, Cas10, and Cas7 were present, but the absence of Cas1 and Cas2 suggests an early system with RNA-based immune response capable of cutting (Cas6/Cas3) and binding (CSM/Cas10) RNA but lacking full CRISPR.

These observations provide compelling hints that cellular life was already involved in an evolutionary arms race with viruses on Hadean Earth.

### 6.2 Metabolism

Perhaps it should not be surprising that the LUCA was an anaerobic acetogen (**Moody et al., 2024**), capable of carbon fixation and energy generation through the Wood-Ljungdahl pathway (WLP) (**McGlynn et al., 2022**). Although rooted in the Bacteria domain, not all acetogens are in Bacteria. While it is true that most classical acetogens (e.g. *Acetobacterium woodii* or *Clostridium sp.*) are Bacteria, certain methanogenic Archaea (e.g. *Methanosarcina acetivorans*) also act as acetogens. In the case of LUCA, it possessed almost all genes for archaeal and most bacterial versions of the WLP (PP > 0.7), enabling acetogenic growth and/or carbon fixation (**Drake et al., 2006**). The WLP is one of the most ancient carbon fixation pathways, requiring only H_2_, CO_2_, and transition metal cofactors found commonly in ultramafic rocks (Ni, Fe, Co) (**Ragsdale and Pierce, 2008; Martin and Russell, 2007; Mojzsis, 2026**). This kind of metabolism is completely consistent with a chemolithoautotrophic lifestyle expected to take advantage of Hadean geologic environments, wherein energy is derived from H_2_ oxidation coupled to CO_2_ reduction. Hydrogenases catalyze the reversible oxidation of H_2_, providing electrons for CO_2_ reduction and energy generation. The LUCA encoded NiFe hydrogenase subunits (K06281, PP = 0.90; K14126, PP = 0.92), enabling growth on hydrogen (e.g. **Vignais and Billoud, 2007**).

We have argued that photosynthesis came later. Before the advent of photosynthesis, primary productivity would have been destitute. Global net primary productivity of the early chemoautotrophic biosphere composed of an acetogenic LUCA and (plausibly) methanogens, is estimated at ∼1 × 10¹² to 7 × 10¹² mol C yr⁻¹ (**Canfield, 2005**), which is approximately three orders of magnitude less than that of today (∼10¹⁵ mol C yr⁻¹) (e.g. **Field et al., 1998**). On the other hand, hydrogen fluxes were several times higher on early Earth than today due to greater internal heat, with modern volcanism producing ∼1 × 10¹² mol H₂ yr⁻¹ and serpentinization ∼0.4 × 10¹² mol H₂ yr⁻¹ (**Sleep et al., 2004**). Chemoautotrophic life in the Hadean, including the LUCA, was forced to reside near locally rich sources of hydrogen before net primary productivity increased with the invention of photosynthesis. Indeed, it was only after photosynthesis arrived that a huge pool of organic matter became available to provide an abundant and powerful electron donor that allowed life to pursue a vast panoply of new metabolic styles later in the Archean (**Morrison and Mojzsis, 2021**).

## 7. Kinetic Window Comparisons: Earth versus Other Solar System Bodies

### 7.1 Carbonaceous Asteroids: Brief Biocompatibility in an otherwise Flawed System

Carbonaceous chondrite parent bodies (e.g. C-type asteroids) and distant icy objects such as those found in the Kuiper Belt, have been proposed as potential sites for abiogenesis owing to their content of water ice and organic molecules (e.g. **Simankov, 2008; Desch et al., 2009**). These objects are cold now, but numerical models of thermal evolution indicate that they maintained liquid water for tens of millions of years following accretion and depending on how early they formed. Liquid water first appears at ∼1 Myr when the 273 K isotherm is reached, triggered by radiogenic heating from ²⁶Al decay (e.g. **MacPherson et al., 1995**). For example, 200-km carbonaceous asteroid with 40% initial ice content takes ∼60 Myr to cool completely, with mild temperatures (273–373 K) maintained for ∼24 Myr in the center. The overall cooling timescale for a 100-km asteroid with 20% initial ice is ∼10 Myr (**Abramov and Mojzsis, 2011**). Once liquid water is available inside an asteroid, exothermal hydration reactions (e.g., forsterite + enstatite + water → antigorite, releasing 69 kJ per mole serpentine) (**Helgeson et al., 1984**) cause thermal runaway, sometimes exceeding 500 K in a large fraction of the asteroid (**Abramov and Mojzsis, 2011**). If initial ice content is <20%, almost all water is consumed in hydration reactions shortly after melting, with the duration of conditions suitable for biochemistry essentially zero.

Another obstacle to the origin of life in asteroids that may be less obvious but no less important is their extremely low permeabilities (10⁻¹⁹ to 10⁻¹⁷ m²) **(McSween et al., 2002)**. This means that water is essentially immobile, permitting transport length-scales of only ∼100 µm at most, even at 1 Myr timescales. This is six-orders-of-magnitude lower permeability than earlier models (**McCoy et al. 2006**) implies that any prebiotic chemistry would remain very narrowly confined, preventing the mixing and concentration thought necessary for complex chemical evolution. The geometric mean pore size in primitive carbonaceous chondrites (e.g. Acfer 094) is 5–50 nm, presenting a first-order problem for chemistry to reach the level of complexity to lead to life. The smallest known non-virus living organism (*Nanoarchaeum equitans*) is 400 nm in diameter (**Huber et al., 2002**), eight times larger than the largest pores. Even protocells with minimal complexity would require pores >100 nm (**Szostak et al., 2001**).

Even if local permeability was higher and pore sizes larger, owing perhaps to fracturing and other mechanical damage from impacts, biocompatible conditions within an asteroid are confined to relatively narrow, migrating zones. In a typical asteroid with hydration as a major heat source and short-lived radionuclides operative for <3 Myr after Solar System formation, conditions favorable for prebiotic chemistry are confined to outer regions immediately after thermal runaway (central regions too hot), then migrate inward at 1–10 mm yr⁻¹ as the asteroid cools (**Abramov and Mojzsis, 2011**). The duration of biocompatible conditions at any given location within a carbonaceous asteroid up to 200 km in diameter does not exceed ∼3 Myr, even if central temperatures >273 K persist for >8 Myr.

#### Asteroid kinetic windows

While carbonaceous asteroids maintained liquid water for ∼10–60 Myr, severe constraints from hydration reactions (removing water), low permeability (preventing mixing), and small pore sizes (excluding cells) make them unlikely sites for abiogenesis. Larger asteroids and dwarf planets (Ceres: 950 km diameter; Themis family parent bodies: 390–450 km) are more promising, with predicted liquid water oceans lasting millions of years and possible hydrothermal activity extending for even longer (**McCord and Sotin, 2005; Castillo-Rogez and McCord, 2010**). Despite what may appear to be promising for abiogenesis, biocompatible conditions within asteroids are short-lived, low volume, bereft of reliable disequilibrium chemistry, bathed in radiation and subject to disruption from collisions. For these reasons and more, chemical precursors for life have been found in asteroidal meteorites but no trace of indigenous life has ever been discovered in a carbonaceous chondrite.

### 7.2 Mars: Early Biocompatibility with Lethal Drawbacks

Mars has long been a compelling target for astrobiology due to evidence for ancient liquid water, including valley networks, lake deposits, and hydrated minerals. The martian surface was potentially biocompatible during the Pre-Noachian period (∼4.48-4.1 Ga), overlapping with the abiogenesis window on Earth.

#### 7.2.1 Martian Advantages

Mars had several advantages for an independent abiogenesis over other planets aside from Earth. Valley networks and lake deposits indicate sustained liquid water for >10^7^ years during the (pre-)Noachian (**Jakosky and Phillips, 2001**; **Carr and Head, 2010; Poulet et al., 2005**). Impact-generated and volcanic hydrothermal systems provided chemical disequilibria (**e.g. Abramov and Mojzsis, 2016**). It has been proposed that early Mars hosted a CO_2_-H_2_ atmosphere, favorable for prebiotic synthesis (**Wordsworth et al., 2013**; **Woo et al., 2019**). Finally, Mars’ lower gravity (0.38 g) may have facilitated the formation of deep pore networks in the crust (**Knapmeyer-Endrun et al., 2025**).

#### 7.2.2 Martian Disadvantages

Mars also faced significant limitations to its ability to provide conditions for a sustained origin and development of early life. Mars’ low gravity and lack of a global magnetic field over geologic time led to atmospheric loss, reducing surface pressure and destabilizing liquid water over most of its history (**Brain and Jakosky, 1998; Jakosky et al., 2017**). Even if Mars did possess a thicker early atmosphere, its greater distance from the Sun (∼1.5 AU) required strong greenhouse warming to maintain liquid water that is difficult to reconcile with climate models (**Haberle, 1998; Wordworth et al., 2013**). Climate models suggest that early Mars was cold with transient warming (e.g. **Fastook and Head, 2015**). The martian surface has long been exposed to intense UV owing to a thin atmosphere, requiring subsurface, mineral, water-column (e.g. **Michalski et al., 2018**) or perhaps even pigment shielding to protect lifeforms.

##### Martian Kinetic windows

The kinetic windows for abiogenesis on pre-Noachian Mars were likely shorter and more transient than on Hadean Earth. Liquid water at the surface may have been present only during impact events, massive volcanic eruptions, or extreme orbital variations that temporarily warmed the climate (**Head et al., 1999; Wordworth, 2016**). This transient nature for biocompatibility on pre-Noachian Mars severely curtailed the duration of continuous chemical evolution.

### 7.3 Europa and Enceladus: Promising? Or Gravely Limited?

Jupiter’s moon Europa and Saturn’s moon Enceladus possess subsurface oceans beneath ice shells, maintained by tidal heating and/or fossil heat from formation or tidal evolution (**Pappalardo et al., 1999; Porco et al., 2006**). What is little discussed, but important in the kinetic windows discussion, is how long these cloaked oceans have existed. In the case of some if not all of the Galilean moons – with the exception of Io – (Europa, Ganymede and probably Callisto) it may be that subsurface oceans have existed there for billions of years (**Roberts and Nimmo, 2008**), in which case these provide extended kinetic windows for abiogenesis. Despite much optimism over the “habitable potential” of these icy worlds which have driven missions to explore them for their astrobiological potential, several important parameters militate against abiogenesis having occurred in them (e.g. **Mojzsis, 2021, 2026**).

First of these is the nature of the watery regime that these icy worlds support. One of the principal advantages of Earth over any other world that we know is that it has sustained liquid water on its geologically active surface in contact with atmosphere over billions of years. Tidal dissipation translated to passive heating maintains liquid water in ice-covered oceans like those of Europa for >10⁹ years, far exceeding the abiogenesis window on Earth defined in **Abramov et al.** (**2026**). Other icy moons, such as Ganymede and Callisto (Jupiter) do not sustain liquid water by tidal heating, but instead by intrinsic radioactive decay and leftover gravitational heat. Titan (Saturn) may have patches of liquid water near its core (see below). Suffice to say, even with liquid water present the kinetic barriers to abiogenesis are extreme.

It is possible that disequilibrium chemistry and pH gradients can exist in interaction between the ice shell and the water mantle such as with Europa (**Glein et al., 2015**), but this is unreliable and arguably not enough to sustain a biosphere (**Mojzsis, 2021; cf. Chyba and Hand, 2001**). Plume samples from Enceladus contain organic molecules, including what have been argued to be amino acids and other complex hydrocarbons (**Postberg et al., 2018**). Yet, none of these have been claimed to be proof of life, past biological activity or even of active prebiotic chemistry. The molecules were most likely leached from the carbonaceous building blocks of Enceladus (**cf. McKay et al., 2008**).

While tidal heating manages to keep water liquid beneath ice shells but has never provided as much energy per unit volume as radiogenic heating from a rocky base even in the early days of the Solar System (e.g. **Tobie et al., 2008**). We emphasize the fact that no rocks can melt within any known icy moons, which in turn strictly limits productivity owing to the absence of any meaningful chemical disequilibrium from alteration of fresh igneous rock to support biological functions. Icy moons have low rock/water ratios, limiting the efficacy of serpentinization reactions even if fresh rock was available and mitigating effective leaching of phosphorus and other rock-derived nutrients (**Waite et al., 2009; Glein and Waite, 2020**). In all cases, the ice shells of cloaked oceans of the outer Solar System isolates the ocean from surface zone (either with or without and atmosphere). The surface of even airless icy moons acts as an electron acceptor at the planetary scale, and separation of the cloaked ocean from the surface militates access to atmospheric photochemistry and/or UV-driven synthesis, which severely limits the diversity of prebiotic precursors (**Chyba and Philipps, 2001**). Ocean temperatures in the icy satellites are likely 0-10°C (**Zolotov and Shock, 2001**) slowing reaction rates in an already dilute solution, and thus potentially preventing the emergence of complex chemistry.

#### Icy World Kinetic windows

We are necessarily skeptical of the biocompatibility of cloaked oceans locked within icy worlds around the giant planets, in the outer Solar System, or between the stars – rogue planets – unless they are large; in the range of an Earth mass (**Stevenson, 1999**). The extended duration of liquid water on Europa and Enceladus ostensibly provides long kinetic windows, but the extremely low energy flux and nutrient scarcity may prevent the emergence of life (**Chyba, 2000**). If life ever existed, it likely relied on pseudo-heterotrophy and chemolithoautotrophy (similar to the pre-LUCA biosphere on Earth) before all resources were lost to it as the system reached chemical equilibrium with effective hydrolysis of all rocky material, and the demise of radiogenic heat.

### 7.4 Titan: Ultracold and Exotic Chemistry

Saturn’s moon Titan possesses a thick nitrogen atmosphere, surface lakes of liquid methane and ethane, and a subsurface water ocean (**Lorenz et al., 2006; Lunine and Atreya, 2008**). This unique environment offers the possibility of exotic, non-water-based life (**McKay and Smith, 2005**). Any discussion of the chemistry of life outside of the pressure-temperature-compositional bounds of known terrestrial biochemical processes is pure speculation until we find it.

#### 7.4.1 Titan Advantages

Atmospheric photochemistry on Titan produces complex organic molecules, including amino acids, nucleobases, and polycyclic aromatic hydrocarbons (**Wilson and Atreya, 2004; Hörst et al., 2012**). Methane and ethane lakes provide a liquid medium for chemistry, albeit at cryogenic temperatures (∼90 K) (**Lunine et al., 1983**). Coupled atmospheric photochemistry and subsurface water-ice+rock interactions on and in Titan have the potential to provide chemical disequilibria, but it is not possible to place constraints on this at the present time.

#### 7.4.2 Titan Disadvantages

At 90 K, reaction rates on Titan’s surface are extremely slow (10⁶–10¹² times slower than at 300 K for typical activation energies) (e.g. **Benner et al., 2004**), which extends kinetic windows to >10⁹ years. Methane and ethane are non-polar, preventing the formation of amphiphilic membranes and limiting the types of chemistry possible (**McKay and Smith, 2005; Stevenson et al., 2015**). Subsurface patches of liquid water close to the core are isolated from the surface by an ice shell, preventing interaction with atmospheric organics which mitigates chemical disequilibria.

##### Titan’s Kinetic windows

Titan’s kinetic windows are extremely long due to slow reaction rates, but the feasibility of life in liquid methane remains speculative. If life exists in some environment on Titan, it will require fundamentally different biochemistry to function than what we know of terrestrial life and may even be difficult to detect or recognize.

### 7.5 Earth’s Unique Advantages

Comparing Earth with other Solar System bodies highlights several unique advantages that created favorable kinetic windows for abiogenesis on our planet **(Table 1**):

1. **Sustained liquid water**: Earth has maintained surface liquid water continuously since about 4.4 Ga and perhaps earlier, far longer than asteroids (∼10⁷ years) and more accessible than subsurface oceans on icy moons, or water-rich layers in the atmosphere (Venus? Jupiter?).
2. **Wet-dry cycling**: The presence of exposed landmasses, rainfall, and evaporative ponds on Hadean Earth created ideal settings for wet-dry cycling, which is required for nucleotide synthesis and polymerization.
3. **Moderate temperatures**: Earth’s surface temperatures (0–100°C) fall within the optimal range for RNA stability and ribozyme activity, unlike the suboptimal temperature fields in asteroids (thermal runaway to >500 K) or icy moons (-183 – +10°C).
4. **Atmospheric photochemistry**: Earth’s atmosphere provided a source of reduced C-N species (HCN, formaldehyde) through photochemistry and lightning (**Chyba and Sagan, 1992**), while UV radiation was moderated by the water column, minerals and organic hazes.
5. **Spatial scales**: Earth’s large size (5.1 x 10^6^ km^2^ surface area) provided diverse environments (hydrothermal vents, ponds, ocean, atmosphere) connected by hydrological and atmospheric cycles, allowing migration between synthesis and stability environments. Asteroids are too small (100–200 km) for such diversity, while icy moons are isolated by ice shells and water-rich layers in an atmosphere are ephemeral.

**Table 1.** Comparative kinetic window parameters for abiogenesis across planetary bodies and selected small bodies in the Solar System. Values represent estimates from geochemical modeling, meteorite analysis, and spacecraft data; ranges reflect current uncertainties. T = temperature; [P] = bioavailable phosphorus concentration; aw = water activity; Δt = estimated duration of the habitable/kinetic window. See text for references.

| Parameter | **Early Earth (Hadean–Eoarchean)** | **Early Mars (Pre-Noachian – Noachian)** | **Carbonaceous Chondrite Parent Bodies** | **Europa (present)** | **Enceladus (present)** |
|---|---|---|---|---|---|
| **Temperature (T, °C)** | 20–100 (peak ~70; modern ~15) | −10 to 50 (early warm-wet phase) | 0–150 (aqueous alteration) | 0–50 (hydrothermal vent plumes) | 50–200 (hydrothermal chimneys) |
| **Optimal T for RNA (°C)** | 40–70 | Marginal (≤30) | 20–80 (brief) | ~50 (vent-proximal) | ~80 (vent-proximal) |
| **Ocean/fluid pH** | ~5.0–7.5 (acidic → neutral) | 3.0–8.0 (highly variable) | 6.5–9.0 (carbonate-buffered) | 8.0–11.0 (alkaline) | ~8.5–10.5 (alkaline) |
| **Optimal pH for RNA (units)** | 6.0–7.0 | Marginal (acidic phases) | 6.5–8.0 | Marginal (too alkaline) | Marginal (too alkaline) |
| **Bioavailable [P] (µM)** | 1–100 (meteoritic schreibersite) | ~10–500 (evaporite enrichment) | 100–10,000 (schreibersite-rich) | ~0.01–10 (ocean estimate) | ~0.1–10 (plume measurement) |
| **Water activity (aw)** | ~0.98–1.00 (open ocean) | 0.5–0.95 (episodic surface water) | 0.85–0.99 (during alteration) | ~0.98–1.00 (sub-ice ocean) | ~0.98–1.00 (sub-ice ocean) |
| **Wet-dry cycling** | Yes (tidal flats, hydrothermal) | Yes (episodic; aeolian desiccation) | Limited (transient; impact melt) | No (permanently submerged) | No (permanently submerged) |
| **UV flux (W m<sup>−2</sup>)** | ~100–1,000 (early; high EUV) | ~200–800 (after 4.2 Ga; thin atm.) | Negligible (deep body interior) | Negligible (sub-ice; shielded) | Negligible (sub-ice; shielded) |
| **Duration of kinetic window (Myr)** | **200–800**<br>(constrained by LUCA ~4.2 Ga) | ~50–500 (early wet epoch) | **~10–100** (aqueous alteration phase) | Ongoing (>1 Gyr persistent) | Ongoing (>1 Gyr persistent) |
| **Kinetic window overlap score** | **High**<br>(all parameters simultaneously favorable) | **Moderate**<br>(T, [P] favorable; pH, wet-dry marginal) | **Low–Moderate**<br>(T, [P] favorable; no wet-dry; brief) | **Low**<br>(T, aw favorable; pH, wet-dry, UV unfavorable) | **Low–Moderate**<br>(T, aw, [P] moderate; pH, wet-dry unfavorable) |
| **Abiogenesis plausibility** | **Demonstrated** | Possible (early epoch) | Unlikely (duration, no wet-dry) | Unlikely (pH, wet-dry) | Possible (chemical richness) |

*Notes:* Phosphorus concentrations for modern Earth ocean ~2 µM; carbonaceous chondrite parent body estimates from (**Abramov and Mojzsis, 2011**); Europa ocean pH from Galileo magnetometry and geochemical models; Enceladus plume composition from Cassini INMS (**Hansen et al., 2020**). Kinetic window overlap score is a qualitative synthesis of all parameters listed.

These advantages created overlapping kinetic windows spanning several hundred million years, allowing the sequential stages of abiogenesis (precursor synthesis, polymerization, functional emergence, cellular organization, evolutionary innovation) to occur in a coordinated manner.

## 8. Discussion

### 8.1 Why the RNA World Was Brief

Our central finding is that the RNA World was a brief, transient stage which lasted less than 130 Myr in the early part of the Hadean eon. This compressed timeline reflects fundamental kinetic constraints imposed by RNA’s chemical instability, particularly its susceptibility to alkaline hydrolysis and inability to form stable hydrophobic cores. This instability provides physical limitations, but limits to complexity and function inherent to an RNA or RNP organism means that these kinds of life are evolutionary dead ends; we propose that life would have gone extinct if DNA did not arise in response to global changes to conditions during the early geological evolution of Earth.

The kinetic drive the led ultimately to the DNA transition arises from two factors:

1. **Complexity limits**: The high error rate (∼10⁻² to 10⁻³ per nucleotide) of RNA (**Bartel Szostak, 1993**) limits genome size to ∼100–1000 nucleotides (**Biebricher and M. Eigen, 2006**), severely constraining metabolic and regulatory complexity. The LUCA genome was ∼2.5 Mb encoding ∼2,600 proteins; this is three orders of magnitude larger than a postulated RNA organism genome, implying that the transition to DNA (with its lower error rate) was essential for genomic expansion.
2. **Functional limitations**: Polyanionic charges in the structure of RNA prevent formation of hydrophobic cores and membrane integration (**Von Heijne et al., 2006**), limiting the range of catalytic reactions and cellular functions accessible to ribozymes (**Cech, 2000**). The incorporation and evolution of proteins, which can form hydrophobic cores and integrate into membranes, was necessary to overcome these limitations for a time. But, without DNA, an RNP organism would not be able to adapt new life strategies such as ever more elaborate metabolic styles and trophic community structure. It simply did not have the capacity to do this.

The transition from RNA to DNA via RNP was thus not merely an evolutionary refinement but a kinetic necessity, driven by environmental pressures that made RNA-based life increasingly untenable. The fact that DNA came after proteins (rather than proteins after DNA) is evidenced by the absence of DNAzymes with charge-neutralized backbones: if DNA had evolved first, the “shorter evolutionary distance” to invent such DNAzymes would likely have been traversed (e.g. **Forster and Walde, 2022**).

### 8.2 LUCA as a Survivor, not a Pioneer

The assigned age of the emergence of the LUCA (4.09–4.33 Ga) and complexity (Bacteria-grade) (**Moody et al., 2024**) shows that it was not the first organism but a survivor of earlier evolutionary diversification (**Woese, 2002**). Furthermore, the LUCA was part of an established ecological system, with its acetogenic metabolism requiring either co-existing autotrophs to provide organic compounds (if organoheterotrophic) or creating niches for other metabolisms through its by-products (if chemoautotrophic) (**Ferry and House, 2006**). The story of the evolution of life from the very beginning is defined by the survival of those just good enough, who lived long enough, to reproduce.

This ecosystem context has several important implications for our understanding of the Hadean biome.

Foremost, is the existence of an ecosystem in the mid-Hadean implies – but does not yet prove – that life had already diversified into multiple metabolic guilds (autotrophs, heterotrophs, possibly methanogens) (e.g. **Martin and Russell, 2007**). The bottom line is that LUCA represents the last common ancestor of all *extant* life, but earlier lineages – that would have been the LUCA but are not – may have existed and gone extinct. Furthermore, the LUCA’s early CRISPR-Cas system indicates that viruses were already present and exerting selective pressure. This observation supports the view that the virus-host arms race is as ancient as cellular life itself (**Koonin et al., 2006; Forterre, 2006**). Lastly, this early evolution was rapid. The short interval from the onset of habitability (4.33 Ga) to LUCA (**Abramov et al., 2026**) implies rapid evolutionary innovation, consistent with the faster evolutionary tempo of RNA- and RNP-based systems (**Medvegy et al. submitted**). The transition to DNA genomes after LUCA dramatically slowed this tempo by increasing replication fidelity.

### 8.3 Kinetic Accessibility: A Quantitative Framework

The preceding discussion of wet-dry cycling highlights a general challenge in origin-of-life research: while environmental conditions may be thermodynamically permissive, the question of whether a critical reaction will actually occur within available geological time is fundamentally kinetic. Here we formalize this distinction through what we describe as a simplified albeit quantitative kinetic accessibility framework that integrates rate constants, reactant concentrations, reactive volumes, cyclic frequencies, and temporal windows into a single evaluative inequality.

#### 8.3.1 The Master Inequality

Borrowing from nuclear physics and collision chemistry, the expected number (*N*) of productive molecular events for a given prebiotic reaction can be expressed as:

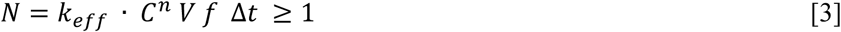

where *k_eff_* is the effective rate constant under local conditions (units: *M*^1−*n*^*s*^−1^), *C* is the reactant concentration (*M*, or the product of concentrations for multi-reactant steps), *n* is the reaction order, *V* is the reactive volume (*L*), *f* is the recurrence frequency for cyclic processes (cycles *s*^−1^; *f* = 1 for continuous reactions), and Δ*t* is the duration of favorable conditions (*s*). The left half of the equation *k_eff_* · *C^n^* is built entirely on standard Collision Theory and the Arrhenius Equation. In modern macro-scale synthesis, the adaptation of microscopic kinetics into a macroscopic parameter framework by explicitly multiplying chemical rates by global volume (*V*), time windows (Δ*t*), and survival fractions (*f*) to set a threshold of *N* was popularized by a number of workers (e.g. **Capra and Luisi, 2014**). Suggested values for these are provided in **Table 2**. The criterion *N* ≥ 1 demarcates kinetically accessible pathways from merely thermodynamically possible ones (**Martin and Russell, 2007**).

**Table 2.** Kinetic accessibility variable summary and early Earth ranges.

| **Symbol** | **Definition** | **Typical early Earth value/range** |
|---|---|---|
| *k_eff* | Effective rate constant | 10<sup>-9</sup>-10<sup>-2</sup> s<sup>−1</sup> (uncatalyzed); up to 10<sup>3</sup> M<sup>−1</sup>s<sup>−1</sup> with mineral catalysis |
| *C* | Reactant concentration | 0.2 μM – 10 mM (nucleobases); 10–100 μM (ribose) |
| *n* | Reaction order | 1 (first-order); 2–3 (multi-reactant condensations) |
| *V* | Reactive volume | 10<sup>−18</sup> L (vesicle) → 10<sup>21</sup> L (ocean) |
| *f* | Recurrence frequency | 1 per 15 h (diurnal); 1 per year (seasonal); 0.1–10 per year (pond evaporation) |

For scenarios in which *N* is small (*N* ∼ 0.01–100), a Poisson stochastic interpretation is more appropriate than the deterministic inequality. That is because we wish to model real-world events as a random, continuous-time process where events occur independently at a constant average rate. This is the conceptual framework behind the Poisson process, a cornerstone of probability theory used to simulate randomness over time or space. Hence, the number of events *k* follows a Poisson distribution, which is ideal for modeling productive events such as chemical reactions. The probability that at least one productive event occurs is calculated by finding the probability of the opposite scenario (that zero events occur) and subtracting it from 1. As such, the probability that at least one productive event occurs is:

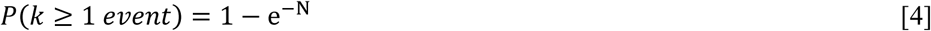

The *N* represents the average expected number of events in a given interval. Here, *e* is Euler’s number (approximately 2.718). Giving *P* ≈ 1% for *N* = 0.01, *P* ≈ 63.2% for *N* = 1 (the feasibility threshold), *P* ≈ 99% for *N* = 5, and *P* ≈ 100% for *N* ≥ 100. This well-known formulation is particularly relevant for evaluating the first formation of a self-replicating molecule, which needs to occur only once somewhere on the planet, and once formed, to persist long enough to reproduce again unimpeded by environmental selection pressures (e.g. **Sharma et al., 2023**).

#### 8.3.2 Temperature Dependence

The strong temperature and activation energy dependence of the rate constant *k_eff_* follows the Arrhenius equation:

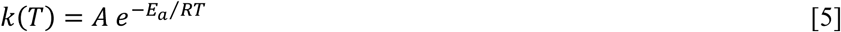

where *A* is the pre-exponential (frequency) factor representing the frequency of collisions with the correct orientation, *E_a_*is the activation energy (kJ mol^-1^), *R* is the gas constant (8.314 J/(mol·K)), and *T* is absolute temperature (K). The exponential sensitivity (**Table 3**) means that plausible early Earth’s ocean temperatures of 350-380 K could increase reaction rates by factors of 10^2^-10^3^× relative to modern ocean temperatures of about 277 K, making pathways accessible within geological windows that would otherwise be kinetically frozen (pun intended) under cooler conditions.

**Table 3.** Activation energy tiers and implied accessibility.

| **Tier** | ***E<sub>a</sub>*** **(kJ mol<sup>−1</sup>)** | **k at 75°C relative to 25°C** | **Representative prebiotic process** |
|---|---|---|---|
| Fast (diffusion-limited) | 0–20 | ~1–3× | Ion pairing, acid-base exchange |
| Moderate | 40–80 | ~10–100× | Nucleotide condensation (mineral-assisted) |
| Slow (uncatalyzed) | 100–150 | ~1,000–10,000× | Peptide bond formation, phosphoester hydrolysis |
| Very slow | >150 | >10,000× | Backbone racemization, high-barrier bond cleavage |

A critical temperature vs. time trade-off follows directly from this analysis. For a target N = 1 representing the feasibility threshold of the prebiotic reaction, the required duration scales as Δ*t* ∝ *e*^−*E*^*^a^*^⁄^*^RT^*. For *E_a_* = 80 kJ mol^-1^, raising *T* from 273 K to 350 K reduces Δ*t* required by a factor of ∼100; raising it further to 375 K reduces it by a factor of ∼1,400. Conversely, reactions requiring 10^8^ years at 273 K could proceed in 10^5^–10^6^ years in a 350-380 K Hadean ocean. Intrinsically, an activation energy of 80 kJ mol^-1^ presents a massive challenge at room temperature (298 K): The term *e*^−*E*^*^a^*^⁄^*^RT^*translates to roughly 1×10^-16^. This means only one out of every ten quadrillion collisions results in a productive event.

#### 8.3.3 Duration (Δ**t**): Temporal Hierarchy of Kinetic Windows

The Δ*t* parameter connects the kinetic framework directly to the geological constraints developed in earlier sections. Different environments on Hadean Earth offered qualitatively different duration windows (**Table 4**). The 200–800 Myr planetary window established in Section 2 is the outer envelope of Δ*t*; individual reactions must achieve N ≥ 1 within the shorter intervals imposed by local environmental stability.

**Table 4.**
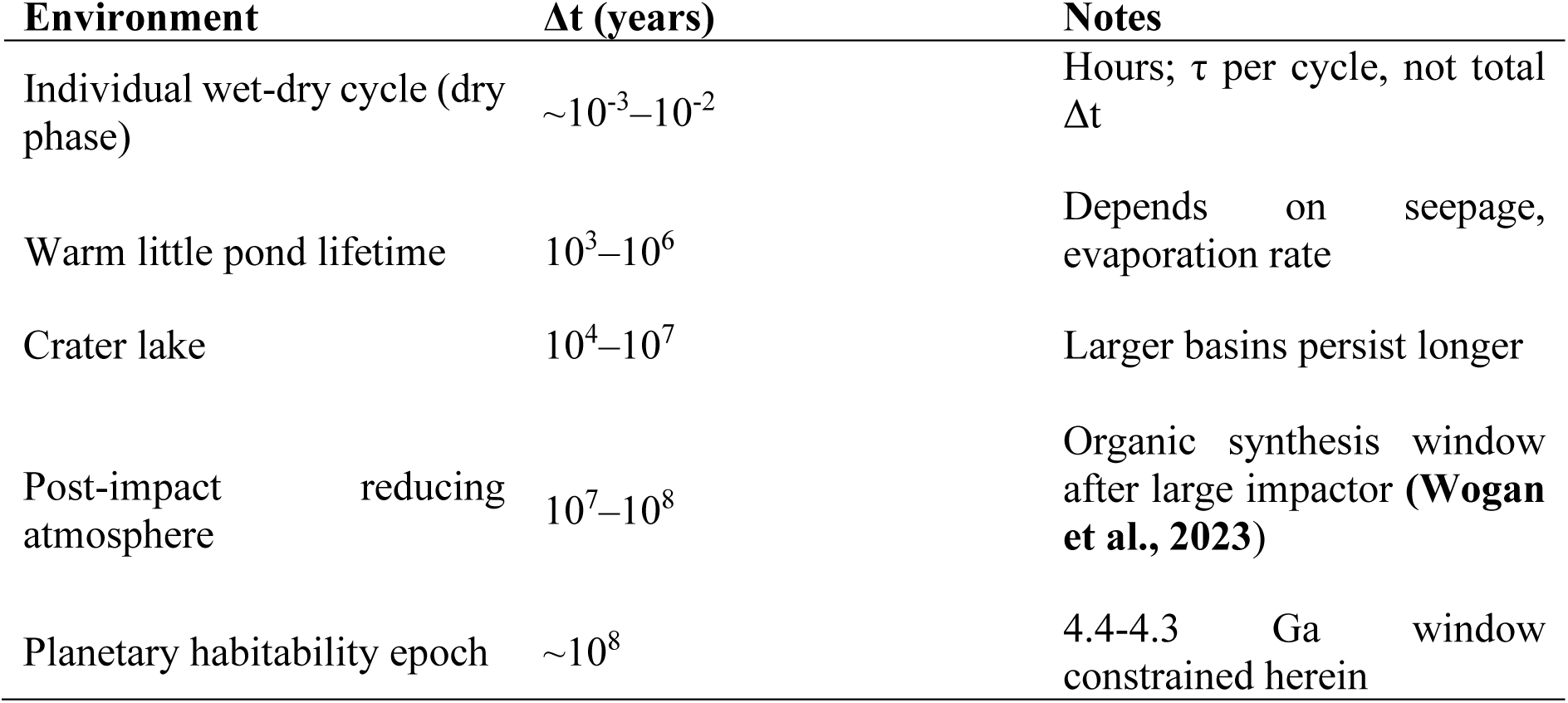
Duration hierarchy for prebiotic kinetic windows.

#### 8.3.4 Spatial Extent (**V**) and the Concentration-Volume Trade-off

For multi-reactant condensations (n ≥ 2), the kinetic accessibility inequality scales as *N* ∝ *V*^(1−*n*)^. Expressing N in terms of total molecular inventory *M_total_* = *C* · *V* :

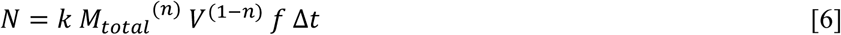

For first-order reactions (*n* = 1), N depends only on *M_total_*, not on V. For second-order reactions (*n* = 2), *N* ∝ *V*^−1^ which means that smaller, more concentrated environments are strongly preferred. For third-order reactions (*n* = 1), *N* ∝ *V*^−2^.

This analysis provides a quantitative explanation for why the warm little (Darwin) pond scenario (**Lane et al., 2010)** outperforms bulk ocean scenarios for multi-reactant prebiotic syntheses: a second-order reaction in a small pond (V = 10^4^ L, C = 1 mM) with the same total molecular inventory as the ocean produces N values ∼10^17^ times larger (**Table 5**).

**Table 5.** Spatial scale tiers and kinetic implications.

| **Environment** | **Typical V** | **C achievable** | **Relative N (n=2) vs. ocean** |
|---|---|---|---|
| Lipid vesicle / protocell | 10<sup>-18</sup>–10<sup>-15</sup> L | mM–M | ~10<sup>36–</sup>10<sup>39</sup>× |
| Mineral pore space | 10<sup>-15</sup>–10<sup>-9</sup> L | mM–M (adsorbed) | ~10<sup>30</sup>–10<sup>36</sup>× |
| Thin film on mineral surface | 10<sup>-9</sup>–10<sup>-6</sup> L | mM–10 mM | ~10<sup>24</sup>–10<sup>30</sup>× |
| Small evaporative pond | 10<sup>3</sup>–10<sup>6</sup> L | 0.1–10 mM | ~10<sup>15</sup>–10<sup>18</sup>× |
| Large lake | 10<sup>9</sup>–10<sup>15</sup> L | µM | ~10<sup>6</sup>–10<sup>12</sup>× |
| Hadean ocean basins | ~10<sup>21</sup> L | nM–µM | 1 (baseline) |

#### 8.3.5 Environmental Parameters

We now explore how the effective rate constant *k_eff_* is modulated by several environmental parameters beyond temperature.

##### Temperature (T)

Optimal range for prebiotic RNA chemistry is 50–100°C, which is hot enough for reasonable *k*, but cool enough for product stability. Above 100°C, RNA hydrolysis rates exceed synthesis rates; below 30°C, activation barriers substantially extend required durations (**Hud et al., 2013**).

##### pH

Nucleotide synthesis is favored at pH 8–10 (nucleophilic attack enhanced at high pH), but RNA stability decreases above pH 7 due to alkaline hydrolysis (*τ*_1/2_ ∝ ^1^⁄_[*OH*_−_]_) (**Forster and Walde, 2022**). The optimal pH window narrows as temperatures increase.

##### Water activity (a_w_)

Condensation reactions (water elimination) are thermodynamically favored at low *a_w_* (dry phase, *a_w_* < 0.5) but require *a_w_* ≈ 1 for reactant diffusion and mixing (**Orgel, 1998**). Wet-dry cycling exploits this by alternating between high-*a_w_* (dissolution/mixing) and low-*a_w_* (condensation) phases.

##### UV flux (Φ)

Absence of an ozone layer and an active young Sun in the Hadean meant that the surface maintained UV fluxes ∼10^2^-10^3^× modern values. Ultraviolet light drives photochemical precursor synthesis (HCN, formaldehyde) but also degrades polymers; mineral shielding or water depth (>1 m attenuates >99% of UV) was likely required for product accumulation.

##### Elemental concentrations [X]

Phosphorus bioavailability, which is critical for nucleotide assembly, required [P] > 10 μM (**Pasek and Kee, 2011**); reduced meteoritic phosphorus species (phosphite, hypophosphite) provided the most bioavailable form the earliest and most intense epoch of late accretion (∼4.5–4.4 Ga) (**Abramov et al., 2026**). Transition metals (Fe, Ni, V, Co) enriched in early oceans (**Mojzsis, 2026**) served as Lewis acid catalysts, lowering *E_a_* for condensation and phosphorylation reactions. As previously mentioned, divalent magnesium is important for RNA catalysis, but it also catalyzes RNA degradation at high pH. Optimal Mg²⁺ concentration at pH 7 is 1– 10 mM, whereas at pH 9, 1 mM Mg²⁺ is deleterious.

#### 8.3.6 Recurrence Frequency (f) and Cyclic Processes

For condensation reactions that are thermodynamically unfavorable in bulk water (ΔG > 0), wet-dry cycling provides kinetic rescue by alternating thermodynamic sign. We have already demonstrated that wet-dry cycling is justified for Hadean Earth with dry land (sec. 2.1). The modified inequality for cyclic processes is:

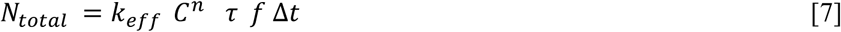

where τ is the duration of the favorable (dry) phase within each cycle. Key frequency ranges on early Earth include diurnal cycles (f ∼ 580 yr^-1^ for a theoretical ∼15-hour Hadean day), seasonal cycles (f ∼ 1 yr^-1^), tidal cycles (f ∼ 365–730 yr^-1^ in coastal zones), and pond evaporation-refilling (f ∼ 0.1–10 yr^-1^) (**Table 6**).

**Table 6.** Cyclic process frequencies and cumulative accessible time.

| **Cycle type** | **f (yr<sup>−1</sup>)** | **τ per cycle** | **Total favorable time over 10<sup>4</sup> yr** |
|---|---|---|---|
| Diurnal (Hadean day) | ~580 | ~7 h dry | ~4,700 yr |
| Tidal coastal pool | ~365–730 | ~6 h dry | ~2,000–4,000 yr |
| Seasonal evaporation | ~1 | ~3 months | ~2,500 yr |
| Pond evaporation-refilling | 0.1–10 | days–months | ~50–10,000 yr |

**Table 7.** Sequential kinetic windows for abiogenesis leading to the LUCA. This compilation incorporates geochemical environments (alkaline crater lakes, warm little ponds, hydrothermal systems) and alignment with different life definitions (Gánti Chemoton and NASA working definition).

| **Evolutional stage (I-V)/ Kinetic window** | **Estimated Timeframe & Duration (Δt)** | **Primary Chemical & Physical Processes** | **Key Environmental Drivers, Settings & Catalysts** | **Rate-Limiting Bottlenecks & Kinetic Constraints** | **Alignment with Life Definitions (Gánti / NASA)** |
|---|---|---|---|---|---|
| **I. Precursor Synthesis Window** | ~4.48 – 4.36 Ga (Δt ~ 10<sup>7</sup>–10<sup>8</sup> yr) | Abiotic synthesis of simple organic precursors: cyanocarbons (HCN), nucleobases, amino acids, ribose, and amphiphiles. Borate-mediated carbohydrate stabilization. | Photochemical reactions in impact-generated reducing (H<sub>2</sub>/HCN-rich) atmospheres; meteoritic schreibersite supplying bioavailable phosphorus (P); alkaline crater lakes / soda lakes (pH 8–10) concentrating borates & carbonates. | High UV photodegradation vs. synthesis rates; low steady-state concentrations of volatile intermediates; alkaline hydrolysis of fragile sugars if not complexed. | Pre-life chemistry; generates fundamental building blocks without molecular memory, metabolic coupling, or heredity. |
| **II. Polymerization & Pre-RNA Window** | ~4.36 – 4.33 Ga (Δt ~ 10<sup>4</sup>–10<sup>6</sup> yr) | Condensation of monomers into functional oligomers; potential ancestral Pre-RNA backbones (PNA, TNA) bypassing pentose instability. | Evaporative wet-dry cycling in terrestrial warm little ponds and alkaline crater lake margins; surface catalysis on volcanic glass (basaltic) and clay minerals. | Dehydration Paradox: Polymerization requires water elimination (*a*<sub>w</sub> < 0.5); ribose chemical instability (t½ < days at high T/pH); high activation barriers (Ea ≈ 100 kJ/mol); spatial trade-off requiring transfer from alkaline synthesis basins to buffered neutral ponds. | Initial assembly of linear informational polymers; early prebiotic 'kinetic bridge' enabling sequence selection. |
| **III. Functional Emergence (RNA World)** | ~4.33 – 4.25 Ga (Δt < 100 Myr) | Emergence of catalytic RNA (ribozymes), template-directed self-replication, and primitive pseudo-heterotrophic catalytic feedback. | Mild freshwater ponds (T ≈ 0–50°C, pH 6–7); low Mg<sup>2+</sup> (1–10 mM) minimizing catalytic self-cleavage; microfluidic mineral pores. | Alkaline Hydrolysis: Rapid phosphodiester cleavage at pH > 7; Eigen's Error Catastrophe: Replication error rate (ε ≈ 10<sup>−2</sup>–10<sup>−3</sup>) caps genome size at 100–1000 nt. | Fulfills NASA criterion for primitive Darwinian evolution; corresponds to Gánti's genetic subsystem (lacks autonomous metabolism and membrane). |
| **IV. Cellular Organization (RNP World)** | ~4.25 – 4.20 Ga (Δt ~ 10<sup>5</sup>–10<sup>7</sup> yr) | Co-evolution of catalytic RNA with cationic/hydrophobic peptides; encapsulation within amphiphilic lipid protocells; proto-ribosome assembly. | Amphiphilic lipid self-assembly in freshwater/brackish ponds; peptide-driven charge neutralization of RNA backbones; geothermal pore networks. | Parasite sequence dominance in bulk solution (mitigated by protocellular spatial isolation); kinetic friction of peptide-ribozyme coordination; membrane permeability trade-offs. | Fulfills Gánti's Chemoton Triad: Full integration of genetic template, primitive metabolic cycle, and protective membrane boundary. |
| **V. Evolutionary Innovation (DNA Transition & LUCA)** | ~4.20 Ga | Transition from RNA/RNP to double-stranded DNA genomes; high-fidelity replication; autotrophic acetogenesis via Wood-Ljungdahl Pathway (WLP). | Hydrothermal/groundwater geochemical disequilibria (H<sub>2</sub>/CO<sub>2</sub> flux); NiFe hydrogenases, ATP synthases, and early CRISPR-Cas defense systems. | Kinetic necessity to escape extinction driven by RNA chemical instability and genome size capping; required evolution of high-fidelity DNA polymerases and repair enzymes. | Fulfills NASA definition: Fully self-sustaining chemical system capable of open-ended Darwinian evolution; Bacteria-grade organism (~2.5 Mb genome, ~2,600 proteins). |

*Notes:* Sequential kinetic windows governing prebiotic evolution on Hadean Earth from initial precursor synthesis to the emergence of the Last Universal Common Ancestor (LUCA) at ~4.2 Ga. Each window represents a temporal interval (Δt) defined by specific physical-chemical parameter spaces (temperature, pH, water activity aw, ionic strength) and rate-limiting kinetic bottlenecks.
Geochemical Setting Integration: Alkaline crater lakes and soda lakes (pH 8–10) play a crucial role during Windows 1 and 2 in **Figure 1** by concentrating borate minerals that selectively stabilize D-ribose against thermal decomposition, and by accumulating high concentrations of cyanocarbon precursors (HCN, cyanamide) in evaporative, closed-basin settings. However, due to rapid alkaline hydrolysis of phosphodiester bonds at pH > 7, long-term functional emergence (**Figure** 1, Window 3) required spatial or temporal transfer into mild, neutral freshwater ponds (pH 6–7) or microfluidic mineral networks, highlighting a fundamental kinetic trade-off between precursor synthesis and informational polymer longevity.
Life Definition Alignment: The transition from a transient RNA/RNP World to a DNA-based cellular architecture was driven by kinetic necessity (avoiding Eigen's Error Catastrophe threshold), enabling the full integration of Tibor Gánti's Chemoton triad (autocatalytic metabolism, genetic polymer, and protective membrane boundary) with NASA's working definition of life as a self-sustaining chemical system capable of open-ended Darwinian evolution.

Even if *N* per cycle ≪ 1, the cumulative *N_total_*= *N_per cycle_*× (f · Δt) can greatly exceed 1. For example, if a reaction yields *N_per cycle_* = 10^-5^ in a single dry phase but cycles operate at f = 365 yr^-1^ over Δt = 10^4^ yr, *N_total_*= 10^-5^ × 3.65 × 10^6^ = 36.5. When *N* values are well above 1, the reactions are favored (highly accessible). The output of this analysis is presented in **Figure 3**.

**FIGURE 3:**
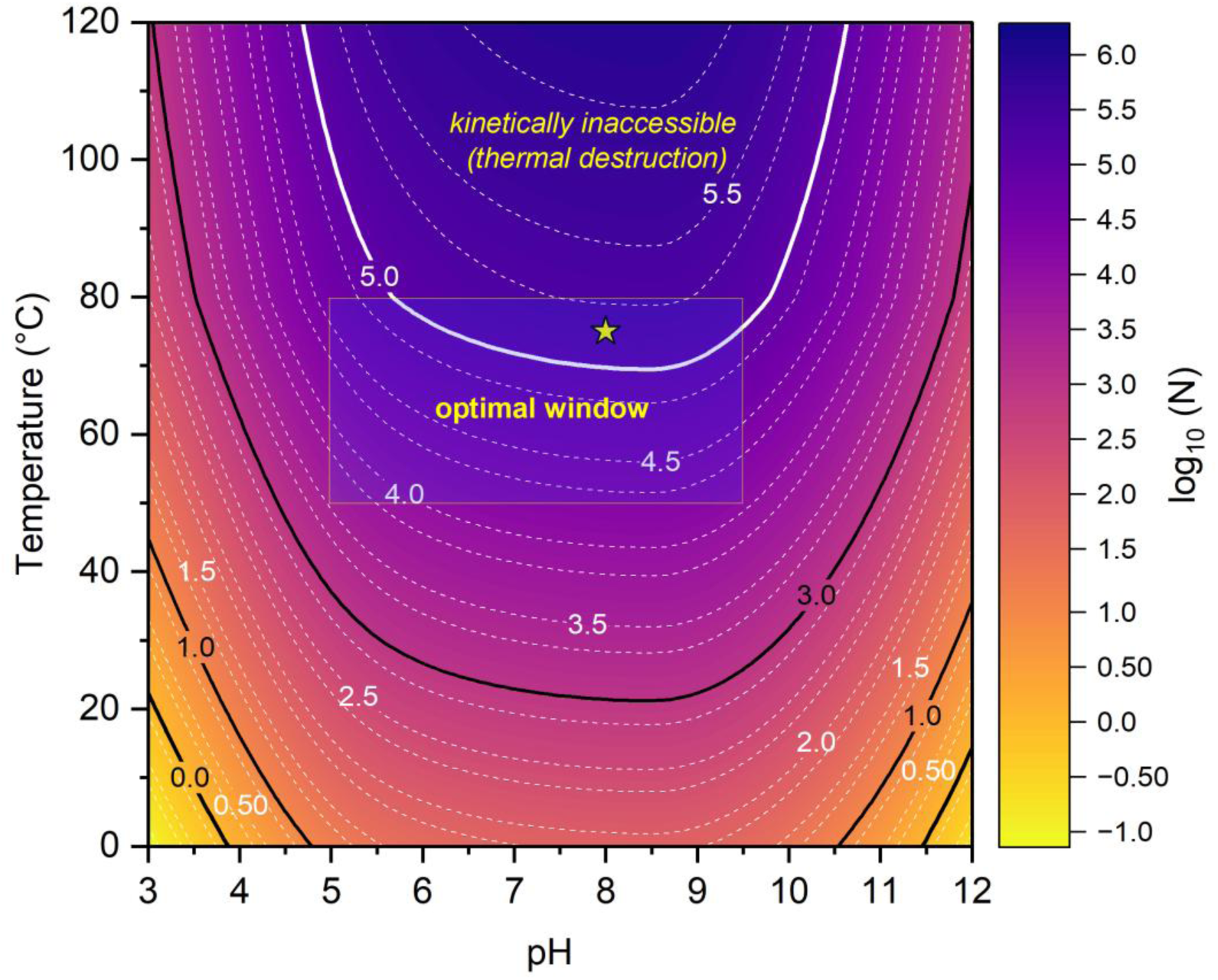
Kinetic accessibility. Kinetic accessibility of nucleotide synthesis across temperature-pH parameter space. Filled contour map of the dimensionless kinetic accessibility index N (mol of nucleotide product) calculated over a grid spanning 0-120 °C and pH 3–12. The color scale encodes log_10_ (N) < 0 where N≪1 (kinetically inaccessible) through to log_10_ (N) ≥ 0 (bold black line at 0.0) defining the critical accessibility threshold N=1 to log_10_ (N) ≥ 5 (bold white line at 5.0) where N=10_5_ (highly accessible). The rectangle denotes the inferred Hadean optimal window for nucleotide synthesis and accumulation (50-80 °C, pH 5–9.5). The star symbol marks the worked example in the text (Sec. 8.3.7; 75 °C, pH 8; log_10_ (N) ≥ 5.2, N≈1.7×10^5^). The optimal window is defined by the RNA thermal stability limit (80 °C), the alkaline hydrolysis onset (pH 9.5) and acid inhibition limit (pH 5). Inaccessible regions are cold/high-pH spaces rate-limited by insufficient thermal activation, whereas hot/low-pH spaces are product-limited by thermal destruction above the stability cutoff.

#### 8.3.7 Simple Worked Examples

##### Example 1: Nucleotide synthesis in a warm little pond (accessible)

Consider ribonucleotide assembly (ribose + nucleobase + phosphate → nucleotide, n = 3) with hypothetical *E_a_* = 80 kJ mol^-1^ and A = 10^10^ s^-1^ at T = 348 K (75°C):

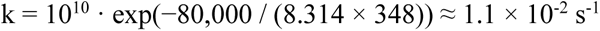

With C_base = 1 mM, C_ribose = 50 μM, C_phos = 10 mM (assumed Hadean pond values), V = 10^4^ L, and Δt = 10^5^ yr = 3.15 × 10^12^ s:

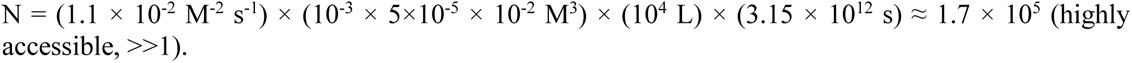

This result vastly exceeds N = 1, confirming that nucleotide synthesis is kinetically accessible in warm, Phosphate-enriched Hadean ponds (**Ritson et al., 2020**) and is consistent with the early planetary window of 4.4 Ga identified in Section 2. The reaction remains accessible (N > 1) even if temperature drops to 25°C, ribose concentration falls to 1 μM, or pond lifetime shortens to 10^3^ yr.

##### Example 2: Peptide bond formation via wet-dry cycling (inaccessible without catalysis)

For uncatalyzed peptide bond formation (n = 2) with *E_a_*= 100 kJ mol^-1^, A = 10^8^ s^-1^, T = 323 K (50°C):

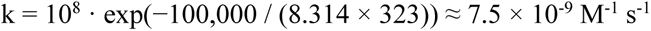

With C = 0.1 M amino acids (evaporation-concentrated), V = 10^-3^ L (thin film), τ = 10^4^ s (3 h dry phase), f = 365 d^-1^ yr^-1^, Δt = 10^4^ yr:

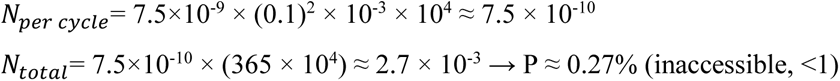

The result looks unsatisfactory. However, if a mineral or glass surface reduces *E_a_* by 30 kJ mol^-1^ (to 70 kJ mol-1), which is achievable, *k* increases by a factor of exp(30,000 / (8.314 × 323)) ≈ 7.3 × 10^4^, yielding *N_total_* ≈ 197 (highly accessible, >1). This calculation quantitatively demonstrates why mineral-assisted wet-dry cycling is a leading scenario for peptide bond formation and why mineral-or glass-catalyzed oligopeptide synthesis (**Jerome et al., 2022; Šponer et al., 2023**) represents a kinetically privileged pathway rather than merely a thermodynamically favorable one.

#### 8.3.8 Planetary-Scale Probability Aggregation

Even when *N* < 1 in any individual environment (*N_single_*), the planetary probability of the event having occurred at least once is dictated by the enormous spatial scale available on Earth:

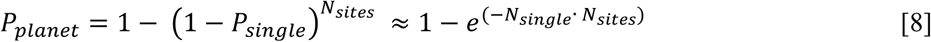

where *N_sites_* is the number of independent environments globally. For *N_single_*= 0.01 and *N_sites_*= 10^6^ warm little ponds (a conservative estimate for a geologically active Hadean Earth with continental nuclei, e.g. **Korenaga, 2021**):

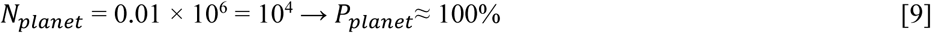

This aggregation logic experiment explains how events that appear statistically unlikely in any single locality, such as the spontaneous assembly of a functional ribozyme, become essentially inevitable at planetary scales over the ∼130 Myr abiogenesis window which we have defined (**Abramov et al., 2026**). Collectively, the kinetic accessibility framework provides a quantitative bridge between the environmental constraints documented here (**Figure 4)**.

**FIGURE 4:**
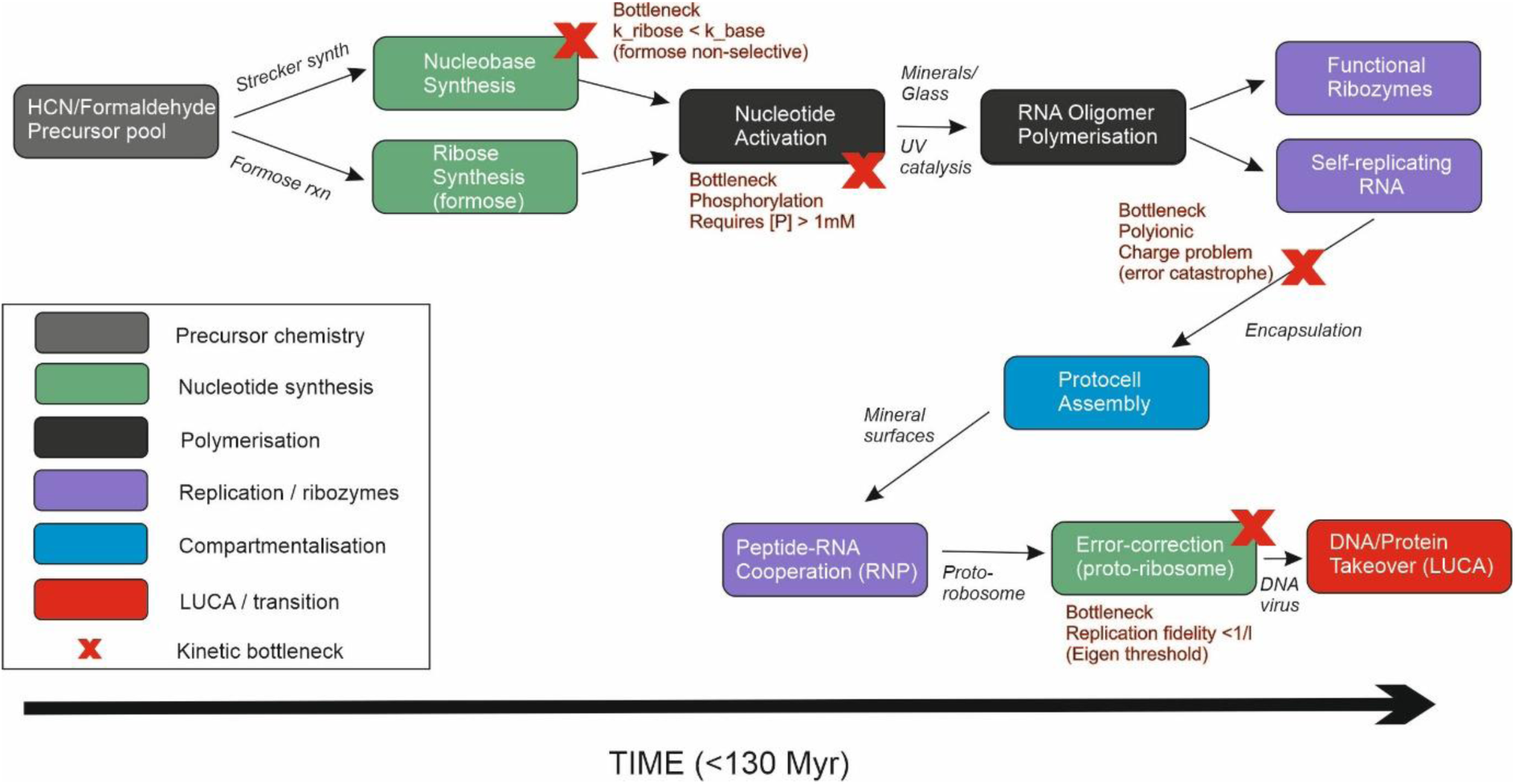
Kinetic Bottlenecks in the RNA World. Schematic diagram illustrating kinetic bottlenecks in the RNA World. Central panel shows the RNA World cycle: nucleotide synthesis → polymerization → ribozyme function → replication → degradation. Each step is annotated with rate constants and half-lives from experimental data. Bottlenecks highlighted in red: (1) Ribose synthesis and stability (t₁/₂ ∼ days to weeks at 25–50°C), requiring borate stabilization or rapid turnover; (2) Nucleotide phosphorylation (ΔG ∼ +20 kJ/mol in water), requiring wet-dry cycling or activation chemistry; (3) Template-directed polymerization (rate ∼ 0.1–1 nt/min), competing with degradation (rate ∼ 10⁻⁶ to 10⁻⁴ min⁻¹ depending on pH and T); (4) Error catastrophe (error rate ∼ 10⁻² to 10⁻³ per nt), limiting genome size to ∼100–1000 nt; (5) Functional sequence discovery (frequency ∼ 10⁻¹¹ to 10⁻¹³ for simple catalysis), requiring stepwise evolution.

### 8.4 Implications of the Kinetic Windows Model for Astrobiology

The framework developed here has several implications for the search for life elsewhere in our Solar System and beyond, to exoplanets. We have emphasized that abiogenesis requires not merely the presence of organic precursors, chemical disequilibria and liquid water, but a temporal coordination of environmental conditions during a planet’s formative period. To foster a biological system from inception, planets must provide overlapping kinetic windows for precursor synthesis, polymerization, functional emergence, cellular organization, and evolutionary innovation. This coordination is more likely on planets with sustained liquid water, wet-dry cycling, and moderate temperatures. Large rocky planets with fluid envelopes – perhaps up to Super Earth in mass (2-6 M_E_) – with diverse environments (hydrothermal vents, ponds, oceans, atmospheres) connected by hydrological and atmospheric cycles provide more opportunities for abiogenesis than small bodies (asteroids), isolated environments (subsurface oceans in icy moons) or geologically moribund worlds (Mars).

These considerations suggest that rocky planets which are broadly Earth-like in mass (>0.5 to <6 M_Earth_) in the habitable zone of Sun-like (G through larger M dwarf) stars with sustained liquid water and effective crustal recycling processes draped in an atmosphere are the most promising targets for detecting life (e.g. **Catling and Kasting, 2017**), as they are most likely to provide the coordinated kinetic windows necessary for abiogenesis. These may be obvious parameters to look for, but they can be difficult to fulfill.

### 8.5 Uncertainties and Future Directions

Despite significant progress in constraining the nature of the Hadean environment over the last two decades, several key uncertainties remain. We do not have a good understanding of Hadean surface conditions (before 4 Ga) beyond the very broad conclusions provided here. The temperature, pH, and atmospheric composition of the Hadean Earth remain poorly constrained due to the scarcity of preserved rocks. Improved models of atmospheric evolution and impact-driven perturbations are needed that are accord with geochemical data. The specific chemical pathways leading from simple precursors (HCN, formaldehyde) to nucleotides and amino acids remain debated. More experimental studies under plausible Hadean conditions (varying pH, temperature, mineral surfaces) are warranted.

While molecular clock analyses appear to place LUCA at in the mid-Hadean, the exact duration of the RNA World and the timing of the RNA-to-DNA transition remain uncertain. Improved calibrations and expanded gene sets may refine these estimates. The extent of ribozyme diversity in the RNA World and the specific catalytic functions they performed remain speculative (**Chen et al., 2004**). Discovery of new ribozymes in modern organisms or through *in vitro* evolution may provide insights.

The mechanisms by which RNA, peptides, and lipids self-organized into protocells with heredity and metabolism is unclear (**Szostak et al., 2001; Luisi et al., 2006**). More experimental studies of compartmentalization and the incorporation of primitive metabolism are sorely needed.

Finally, what of alternative biochemistries? The possibility of alternative genetic polymers (e.g., Peptide Nucleic Acid, PNA; Thiol Nucleic Acid, TNA) or solvents (e.g., liquid methane) remains speculative (**Nielsen, 1993; Eschenmoser, 1999**). Exploring these alternatives further may reveal whether Earth’s biochemistry is unique or representative (**Benner and Sismour, 2005**). We do not know if biochemistry is universal or not.

## 9. Conclusions

Abiogenesis on Earth occurred within a constrained temporal window bounded by planetary cooling to sustain continuously biocompatible conditions (∼4.33 Ga) and the earliest biosignatures (∼3.83 Ga, with contested signals at ∼4.1 Ga). An abiogenesis window of 130 Myr encompassed a immensely complex sequence of prebiotic chemistry, molecular self-organization, and evolutionary innovation, each governed by distinct kinetic constraints.

Key findings include:

1. **Rapid emergence of LUCA**: Molecular clock analyses imply that life evolved to Bacteria-grade complexity soon after continuous habitability. This compressed timeline suggests that once favorable conditions were established, the kinetic barriers to cellular life were surmountable on timescales of 10⁸ years or less.
2. **Brief RNA World**: The RNA World persisted for a time shorter than 130 Myr, terminated by RNA’s intrinsic chemical instability, including its susceptibility to alkaline hydrolysis (half-life inversely proportional to pH above 7) and inability to neutralize polyanionic charges. These and other drawbacks created kinetic bottlenecks that drove the evolution of proteins and DNA.
3. **Critical physical-chemical constraints**: The kinetic windows for abiogenesis were defined by narrow parameter ranges: optimal temperatures of 273–308 K for RNA stability, pH 6–8 for balancing RNA stability against nucleotide synthesis, low Mg²⁺ concentrations (<10 mM) to minimize catalytic degradation, water activity >0.6 for biochemical reactions but <0.9 for polymerization, and phosphorus concentrations >10 µM for nucleotide assembly (**Orgel, 2004**).
4. **Wet-dry cycling as key mechanism**: The presence of exposed landmasses, rainfall, and evaporative ponds on Hadean Earth created settings for wet-dry cycling, which concentrates reactants by 10²–10³-fold, drives thermodynamically unfavorable polymerization reactions, and provides temporal separation between synthesis and function. This mechanism was critical for overcoming kinetic bottlenecks in nucleotide synthesis and RNA polymerization.
5. **LUCA as anaerobic acetogen**: Recent work shows that the LUCA was a Bacteria-grade anaerobic acetogen with the Wood-Ljungdahl pathway, NiFe hydrogenase, ATP synthase, and an early CRISPR-Cas system. It was part of an established ecological system with estimated productivity of ∼10¹²–10¹³ mol C yr⁻¹, three orders of magnitude lower than modern values.
6. **Earth’s unique advantages**: Comparative analysis with carbonaceous asteroids and similar objects (habitable for ∼24 Myr but constrained by 5–50 nm pore sizes and 10⁻¹⁹–10⁻¹⁷ m² permeabilities) highlights the benefits of Earth: sustained liquid water stable at the surface within 150 Myr of planet formation, wet-dry cycling in terrestrial ponds, moderate temperatures, sustained geologic processes (igneous activity), and sufficient spatial scales for chemical evolution.
7. **The RNA/RNP World was bound to fail**: To understand why an RNA World or Ribonucleoprotein (RNP) World was inherently bound for extinction without transitioning to a DNA World, we can look directly through the lens of the prebiotic rate equation discussed previously (sec. 8.3.1): *N* = *k_eff_* · *C^n^ V f* Δ*t* ≥ 1. The biogeodynamic evolution of the Hadean Earth fundamentally degraded the factors of survival (*f*) and time (Δ*t*) for RNA-based life forms. The transition to DNA was not just a minor evolutionary upgrade, rather it was a rescue mechanism to prevent extinction.

In conclusion, life’s origin required not merely the presence of organic precursors but precise temporal coordination of environmental conditions, of kinetic windows for pre-life to evolve to life. Each transition from prebiotic chemistry to RNA, RNA to ribonucleoproteins, and ultimately to DNA-based cellular life, was governed by distinct rate-limiting processes operating over 10⁵–10⁸ year timescales.

Because RNA is chemically unstable, it has a high intrinsic mutation rate. Early ribozymes (RNA enzymes) tasked with copying other RNA molecules lacked sophisticated proofreading mechanisms. Yet, if the mutation rate of a genome is too high, beneficial traits are washed away by a flood of lethal mutations over generations. This is Eigen’s Error Catastrophe. The thermal and chemical stresses of the Hadean Eon meant that early RNA protocells could not expand their genomes beyond a few thousand nucleotides without destroying their own functional codes. They could never evolve complex metabolisms or defenses because they could not store enough stable information. The RNP World evolved from the RNA world to harness amino acids and began building short peptides to act as catalysts and to enhance stability. While this drastically improved the effective reaction rate constant (\[k_{\text{eff}}\]) by providing better enzymes, it worsened the information crisis. The RNP organisms now had to store the code for both them and the supporting proteins. The physical burden of encoding more information broke the constraint of chemical stability such that the RNP world reached a hard bottleneck. The RNP organism desperately needed more complex protein machinery to survive and diversify (especially in metabolic styles), but its unstable RNA genome was physically incapable of safely storing the information to accomplish this.

The kinetic windows model developed here provides a quantitative basis for understanding the probability and timescale of abiogenesis, with implications for the search for life beyond Earth. Planets must provide not only habitable conditions but also the temporal coordination of kinetic windows necessary for the sequential stages of chemical evolution. This coordination is most likely on rocky Earth-like planets in the habitable zone of Sun-like stars. Future research should focus on refining constraints on Hadean surface conditions, elucidating prebiotic synthesis pathways under plausible environmental conditions, discovering new ribozymes that illuminate RNA World diversity, and exploring alternative biochemistries that may reveal whether Earth’s solution to the origin of life is unique or representative.

## Acknowledgements

We thank Oleg Abramov (PSI, BGI), Eörs Szathmáry (Parmenides, ELTE), Steven Benner (FfAME), Elisa Biondi (FfAME), İrep Gözen (Harvard-Radcliffe Institute for Advanced Studies), Thomas Carrell (LMU-München), William “Bill” Martin (Düsseldorf), Dieter Braun (LMU-München), David Catling (UW), Martina Preiner (MPI, Marburg), Michael “Mike” Russell (NASA), John Sutherland (Cambridge), Craig Walton (ETH, Cambridge), Paul Rimmer (Cambridge), Ramon Brasser (CSFK), Leroy “Lee” Cronin (Glasgow) and Ulrich “Uli” Müller (UCSD) for a long and productive history of discussions, debates and insights that contributed to the kinetic windows concept.

## Author contributions

Conceptualization: S.J.M., A.M and B.K. Methodology: S.J.M. Formal analysis: S.J.M. Investigation: A.M., B.K.. Validation: S.J.M., A.M., and B.K. Visualization: S.J.M. Writing—original draft: S.J.M. Writing—review and editing: S.J.M., A.M., and B.K. Resources: S.J.M. Funding acquisition: S.J.M. Project administration: S.J.M. Supervision: S.J.M. and B.K.

## Data and materials availability

All data are available in the main text.

